# Freely-moving mice visually pursue prey using a retinal area with least optic flow

**DOI:** 10.1101/2021.06.15.448520

**Authors:** Carl D. Holmgren, Paul Stahr, Damian J. Wallace, Kay-Michael Voit, Emily J. Matheson, Juergen Sawinski, Giacomo Bassetto, Jason N. D. Kerr

**Author notes:** denotes equal contribution (Carl D. Holmgren, Paul Stahr). Correspondence to: Jason Kerr.

## Abstract

Mice have a large visual field that is constantly stabilized by vestibular ocular reflex driven eye rotations that counter head-rotations. While maintaining their extensive visual coverage is advantageous for predator detection, mice also track and capture prey using vision. However, in the freely moving animal quantifying object location in the field of view is challenging. Here, we developed a method to digitally reconstruct and quantify the visual scene of freely moving mice performing a visually based prey capture task. By isolating the visual sense and combining amouse eye optic model with the head and eye rotations, the detailed reconstruction of the digital environment and retinal features were projected onto the corneal surface for comparison, and updated throughout the behavior. By quantifying the spatial location of objects in the visual scene and their motion throughout the behavior, we show that the image of the prey is maintained within a small area, the functional focus, in the upper-temporal part of the retina. This functional focus coincides with a region of minimal optic flow in the visual field and consequently minimal motion-induced image blur during pursuit, as well as the reported high density-region of Alpha-ON sustained retinal ganglion cells.

## Introduction

The visual system of mice serves a variety of seemingly opposing functions that range from detection of predators, to finding shelter and selection of food and mates, and is required to do so in a diverse set of environments (Boursot, Auffray et al. 1993). For example, foraging in open areas where food is available involves object selection, and in the case of insect predation (Badan 1986, Tann, Singleton et al. 1991), involves prey tracking and capture (Langley 1983, Langley 1984, Langley 1988, Hoy, Yavorska et al. 2016), but the visual system can also simultaneously be relied on for avoidance of predation, particularly from airborne predators (Hughes 1977). Like with many ground-dwelling rodents (Johnson and Gadow 1901) predator detection in mice is served by a panoramic visual field which is achieved by the lateral placement of the eyes in the head (Drager 1978, Hughes 1979, Oommen and Stahl 2008) combined with monocular visual fields of around 200 degrees (Hughes 1979, Drager and Olsen 1980, Sterratt, Lyngholm et al. 2013). In mice, the panoramic visual field extends to cover regions above the animal’s head, below the animals snout and laterally to cover ipsilaterally from behind the animals head to the contralateral side, with the overlapping visual fields from both eyes forming a large binocular region overhead and in front of the animal (Hughes 1977, Sabbah, Gemmer et al. 2017). In addition, eye movements in freely moving mice constantly stabilize the animal’s visual field by counteracting head rotations through the vestibulo-ocular reflex (VOR) (Payne and Raymond 2017, Meyer, Poort et al. 2018, Meyer, O’Keefe et al. 2020, Michaiel, Abe et al. 2020) maintaining the large panoramic overhead view (Wallace, Greenberg et al. 2013) critical for predator detection (Yilmaz and Meister 2013).

Given the VOR stabilized panoramic field of view it is not clear what part of the visual field mice use to detect and track prey (but see: (Johnson, Fitzpatrick et al. 2021). Mouse retina contains retinal ganglion cells (RGCs), the output cells of the retina, with a broad diversity of functional classes (Zhang, Kim et al. 2012, Bleckert, Schwartz et al. 2014, Baden, Berens et al. 2016, Franke, Berens et al. 2017). Given the lateral eye position, the highest overall density faces laterally (Drager and Olsen 1981, Salinas-Navarro, Jimenez-Lopez et al. 2009, Sabbah, Gemmer et al. 2017, Stabio, Sondereker et al. 2018). Further, as the functionally defined ganglion cells (Zhang, Kim et al. 2012, Bleckert, Schwartz et al. 2014, Baden, Berens et al. 2016, Franke, Berens et al. 2017) and cone sub-types (Szel, Rohlich et al. 1992) are segregated into retinal subregions within the large stabilized field of view, recent studies suggest that retinal subregions are tuned for specific behavioral tasks depending on what part of the world they subtend (Hughes 1977, Zhang, Kim et al. 2012, Bleckert, Schwartz et al. 2014, Baden, Berens et al. 2016, Sabbah, Gemmer et al. 2017, Szatko, Korympidou et al. 2020).

The challenge is to measure what part of the visual field the mouse is attending to during a visually based tracking task (Hoy, Yavorska et al. 2016) and the location of all objects within the field of view during the behavior. While recent studies have implied the relationship between prey and retina through tracking head position (Johnson, Fitzpatrick et al. 2021) or measured both the horizontal and vertical eye rotations (Meyer, Poort et al. 2018, Meyer, O’Keefe et al. 2020) during pursuit behavior (Michaiel, Abe et al. 2020) to uncover a large proportion of stabilizing eye-rotations, what is missing is the extent and location of the area used when detecting and pursuing prey, and the relationship to the retina (Bleckert, Schwartz et al. 2014).

Here, we measured the position of a cricket in the visual fields of freely moving mice performing a prey pursuit behavior, using head and eye tracking in all three rotational axes, namely horizontal, vertical and torsional. Eye tracking included an anatomical calibration to accurately account for the anatomical positions of both eyes. To quantify object location in the animal’s field of view and generate optic flow fields, head and eye rotations were combined with a high-resolution digital reconstruction of the arena to form a detailed visual map from the animal’s eye perspective. Given that mice use multisensory strategies during prey pursuit (Langley 1983, Langley 1988, Gire, Kapoor et al. 2016) and can track prey using auditory, visual or olfactory cues (Langley 1983, Langley 1988), we developed a behavioral arena that isolated the visual aspect of the behavior by removing auditory and olfactory directional cues to ensure that the behavior was visually guided. To transfer the retinal topography onto the corneal surface, we developed an eye model capturing the optical properties of the mouse eye. We show that during prey detection mice preferentially position prey objects in stable foci located in the binocular field and undertake direct pursuit. The stabilized functional foci are spatially distinct from the regions of highest total retinal ganglion cell density, which are directed laterally, but coincides with the regions of the visual field where there is minimal optic flow and therefore minimal motion-induced image disturbance during the behavior. Lastly, by building an optical model that allows corneal spatial locations to be projected onto the retina, we suggest that the functional foci correspond to retinal subregions containing a large density of Alpha-ON sustained RGCs that have center-surround receptive fields and project to both superior colliculus and dLGN (Huberman, Manu et al. 2008) and possess properties consistent with the requirements for tracking small and mobile targets (Krieger, Qiao et al. 2017).

## Results

### Forming a view from the animal’s point of view

To measure what part of the visual field mice use during prey capture while also considering that mice can use multisensory strategies during prey pursuit (Langley 1983, Langley 1988, Gire, Kapoor et al. 2016), we first developed an arena which isolated the visual component of prey pursuit by masking olfactory and auditory spatial cues (Figure 1A, see Methods for details). By removing both olfactory and auditory cues, the average time to capture a cricket approximately doubled compared to removal of auditory cues alone (time to capture, median±SD, control 24.92±16.77s, olfactory & auditory cues removed, 43.51±27.82s, p=0.0471, Wilcoxon rank sum test, N=13 control and 12 cue removed trials from N = 5 mice). To track mouse head and eye rotations during prey capture, we further developed a lightweight version of our head mounted oculo-videography and camera-based pose and position tracking system (Wallace, Greenberg et al. 2013) (Figure 1B and Methods, Figure 7 A and B). This approach allowed quantification of head rotations in all three axes of rotation (pitch, roll and yaw), as well as eye rotations in all three ocular rotation axes (torsion, horizontal and vertical, Figure 1C, Figure 1 – figure supplement 1 A and B). The same videography-based system was used to track and triangulate the position of the cricket (see Methods and Figure 1 – figure supplement 1C). To quantify the position and motion of the environment and cricket in the mouse field of view, we also developed a method that enabled a calibrated environment digitization to be projected onto the corneal surface. This approach utilized a combination of laser scanning and photogrammetry, giving a resolution for the reconstruction of the entire experimental room of 2 mm, as well as a detailed measurement of eye and head rotations (Figure 1D-E, and see methods). Mice, like rats (Wallace, Greenberg et al. 2013), have a large visual field of view which extends to also cover the region over the animal’s head (Figure 1F). To ensure the entire visual fields of the mouse could be captured during behavior, we digitized the entire experimental room and contents (Figure 1E, Figure 1 – figure supplement 1D-F, Movie 1). The coordinate systems of the environmental digitization and mouse and cricket tracking systems were registered using 16-20 fiducial markers identified in both the overhead camera images and the digitized environment. The average differences in position of fiducial points between the two coordinate systems were less than 1 mm (mean±SD, x position, 0.18±3.1mm, y position, 0.07±1.6mm, z position, 0.66±1.8mm, N=54 fiducial points from 3 datasets). The next step was to recreate the view for each eye. First, and for each mouse, the positions of both eyes and nostrils were measured with respect to both the head-rotation tracking LEDs and head-mounted cameras, then calibrated into a common coordinate system (Figure 1B). Together, this enabled a rendered representation of the digitized field of view for each combination of head and eye rotations. This rendered image, from the animal’s point of view, contained all the arena and lab objects (Figure 1G-H, Movie 2, Figure 1 – figure supplement 1G). In addition, to object position and distance (Figure 1I), the motion of the environment and each object in the field of view could be quantified as the mouse performed prey capture behaviors (Figure 1J, and Figure 1 – figure supplement 1H).

**Figure 1 with 1 supplement.**
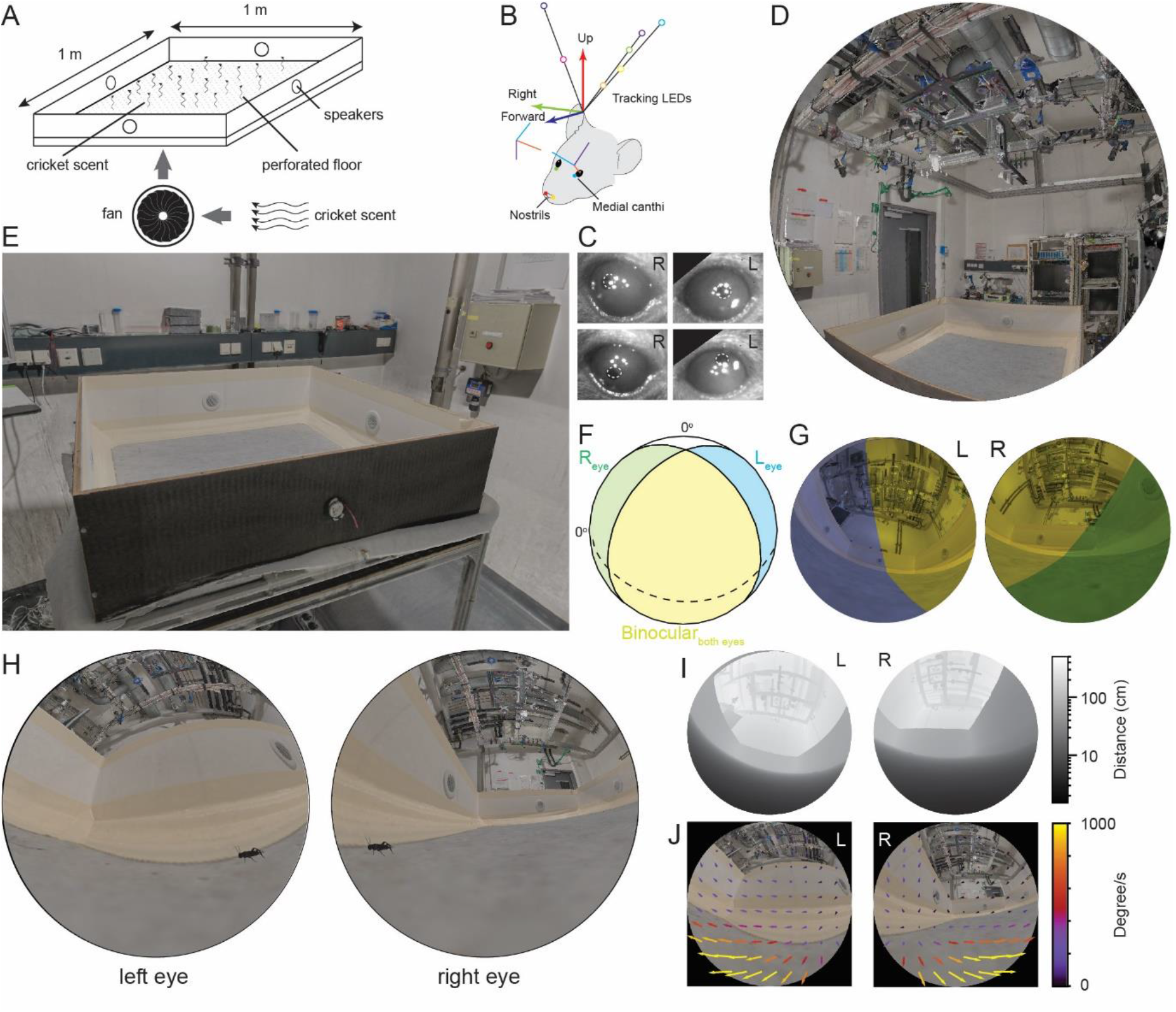
Reconstruction of experimental arena and surrounds from the animal’s perspective. **(A)** Schematic of experimental arena with olfactory and auditory noise. **(B)** Schematic of tracking, anatomical and eye camera calibration. Head position and orientation was tracked using seven IR-LEDs (colored circles). Nostrils (red, yellow filled circles), left (blue filled circle) and right (green filled circle) medial canthi were identified and triangulated in calibration images and used to define a common coordinate system (forward, blue arrow, right, green arrow, and up, red arrow), into which the calibrated eye camera location and orientation could also be placed (eye camera vertical, cyan, horizontal, purple, camera optical axis, red). **(C)** Example left- and right eye camera images with tracked pupil position (white dashed outlines). **(D)** Rendered digital reconstruction of the laboratory room and **(E)** experimental arena. **(F)** Schematic representation of mouse’s left- (blue) and right (green) visual fields, showing also the region of binocular overlap (yellow) and un-seen region (white). **(G)** Reconstruction of the arena and room from the animal’s left- and right eye perspective, with monocular and binocular regions colored as in (F). **(H)** Reconstruction of the animal’s view of the prey (cricket - black) in the experiment arena. (I) Representation of left and right eye views of the arena and surrounding objects grayscale-codedby distance from the eye. **(J)** Rendered animal’s eye views from the left- and right eyes with overlay of arrows representing optic flow during 10 ms during free motion.

### Mice keep prey in a localized visual region during pursuit

Crickets (*Acheta domesticus*), shown previously to be readily pursued and preyed upon by laboratory mice (Hoy, Yavorska et al. 2016), provided a prey target that could successfully evade capture for extended periods of time (total time for each cricket before capture: 64.4±39.3 s, average time±SD, N=21 crickets and 3 mice). To ensure that only data where the mouse was actively engaged in the detection and tracking of the cricket was used, we identified occasions where the mouse either captured the cricket, or contacted the cricket but the cricket escaped (see Methods for definitions), and then quantified the trajectories of both mouse and cricket leading up to the capture or capture-escape (Figure 2A). Within these chase sequences we defined three behavioral epochs (detect, track and capture, Figure 2B, see Methods for definition details) based on the behavior of mouse and cricket, and similar to previous studies (Hoy, Yavorska et al. 2016).

**Figure 2 with 1 supplement.**
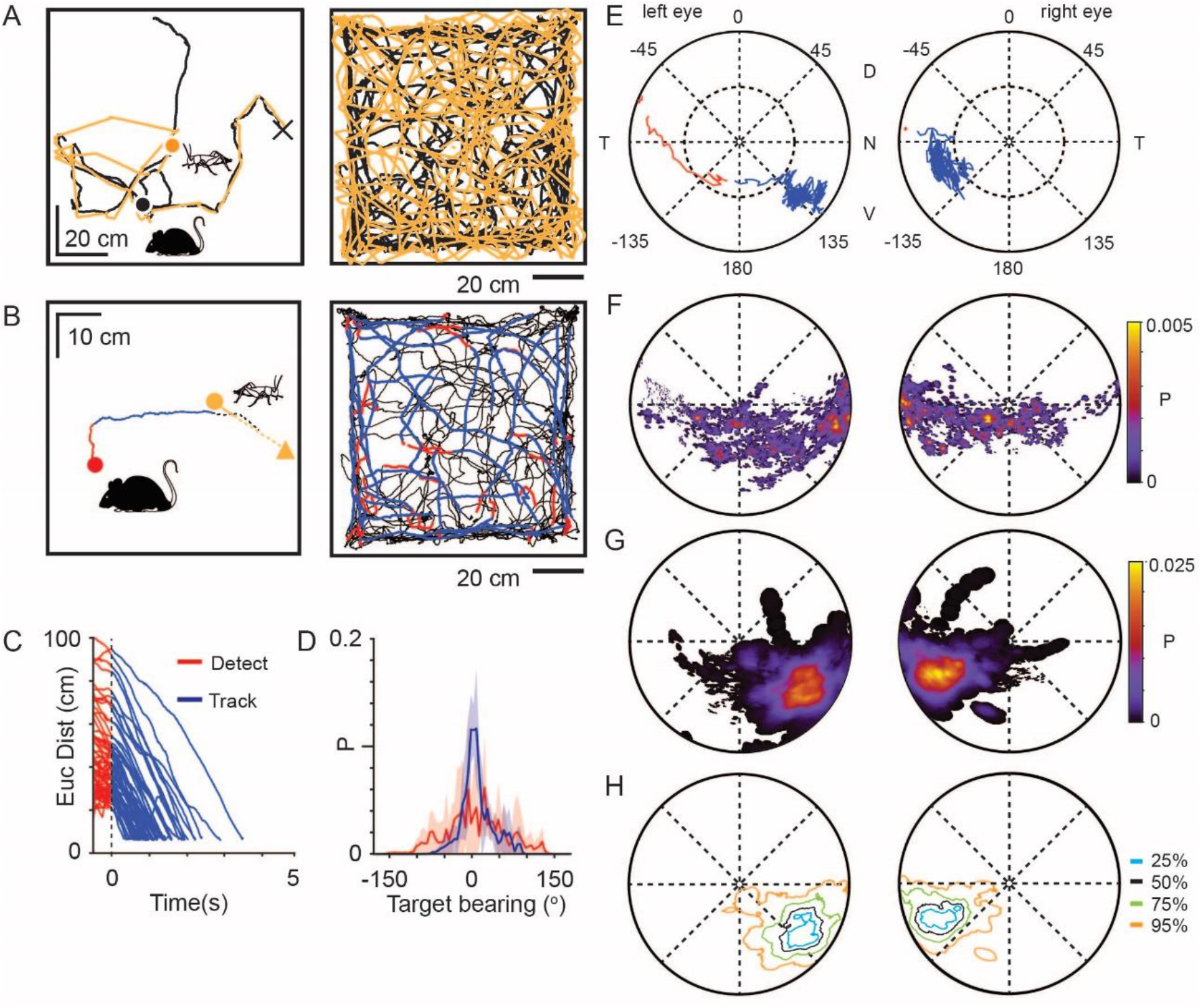
Mice use a focal region of their visual field to track prey. **(A)** Mouse (black) and cricket (orange) paths during a single pursuit sequence (left), and for all pursuit sequences in one session for one animal (right). Pursuit start denoted as filled circles and cricket capture as X. **(B)** Mouse (red and blue) and cricket (orange) paths during an individual pursuit sequence (left) and all pursuit sequences in one session from one animal (right), showing detect (red) and track (blue) epochs of the mouse path. Paths after a cricket escape shown dashed. Pursuit sequence start shown as filled circles, cricket landing point after a jump shown as a filled triangle. **(C)** Euclidean distance between mouse and cricket during detect (red) and track (blue) epochs (n=65 trajectories, n=3 mice). **(D)** Mean and SD bearing to cricket (angle between mouse’s forward direction and cricket location) during detect (red), and track (blue) epochs for all sequences from all animals (detect: 57 epochs, 4406 frames; track: 65 epochs, 13624 frames, n=3 animals, bin size= 5o). **(E)** Trajectory of the projected cricket position in the left and right corneal views, during a single pursuit sequence. Color scheme as for D. The inner dashed circle is 450 from the optical axes. Dorsal (D), ventral (V), nasal (N) and temporal **(T)** directions indicated. **(F)** Average probability density maps for detect epochs (4628 frames from 3 animals). Orientation as in **E. (G)** Average probability density maps for track epochs (13641 frames from 3 animals). Orientation as in E. **(H)** lsodensity contours calculated from the average probability density maps for track epochs. (note that 50% means that this region contains 50% of the total density, and likewise for the other contours). Orientation as in E.

Upon cricket detection, mice oriented and ran towards the cricket, resulting in a significant adjustment to their trajectory (Δ target bearing: 40.2±35.1°, P=6.20X10^−10^, Δ speed: 10.2±7.4cm/s, P=1.91X10^−10^; N=57 detect-track sequences N = 3 mice; Paired Wilcoxon’s signed rank test for both tests), and a rapid reduction in the Euclidean distance to the cricket (Figure 2C). During tracking, the cricket was kept in front of the mouse, resulting in a significant reduction in the spread of target bearings compared to during detect epochs (Figure 2D, Target bearing: detect 6.2± 62.1°, track: 2.5 ±25.6°, mean ±SD, Brown-Forsythe test p=0, *F* statistic=7.05×10^3^, N=4406 detect and 13624 track frames, N=3 mice), consistent with previous findings (Hoy, Yavorska et al. 2016). To avoid the closing phase of the pursuit being associated with whisker strikes (Shang, Liu et al. 2019, Zhao, Chen et al. 2019), tracking periods were only analyzed when the mouse was more than 3 cm from the cricket, based on whisker length (Ibrahim and Wright 1975).

Using the detailed digitization of the behavioral arena and surrounding laboratory method (Figure 1E, Movie 1), an image of the cricket and objects in the environment was calculated for each head and eye position during the predator-prey interaction (Movie 2). Using this approach, we addressed the question of what area of the visual field was the cricket located in during the various behavioral epochs. In the example pursuit sequence in Figure 2E the cricket was initially located in the peripheral visual field and then transitioned to the lower nasal binocular quadrant of the cornea-view during pursuit and capture (red trace in left eye to blue trace in both eyes). Correspondingly, an average probability density map calculated for all animals during the detect epoch showed a very broad distribution of cricket positions across the visual field (Figure 2F, Figure 2 – figure supplement 1A and B). Upon detection the mouse oriented towards the cricket, bringing it towards the lower nasal binocular visual field (Figure 2E). When averaged for all pursuit sequences from all animals, projected cricket positions formed a dense cluster on the cornea of both eyes (Figures 2G and 2H, Figure 2 – figure supplement 1A, C-D, 50% contour center for left and right eye respectively, radial displacement from optical axis 64.3±7.5° and 63.3±9.9°, rotational angle 126.2±8.9° and −115.7±6.1°, mean ± SD, N= 3 mice), which was significantly different from the cluster in the detect epoch (average histogram of the location of cricket image during tracking phase vs average histogram of the location of cricket during detect phase: Left eye P=3.54×10^−46^, Right eye P=1.08×10^−81^, differences calculated by taking the Mean Absolute Difference with bootstrapping, N=57 detect-track sequences, N = 3 mice). Thus, despite mice lacking a retinal fovea (Drager and Olsen 1981, Jeon, Strettoi et al. 1998), the image of the prey is kept on a local and specific retinal area during the tracking and pursuit behavior. The image of the prey was localized on a specific region of retina within the binocular field, while the region of elevated density of RGCs has been found to be located near the optical axis (Drager and Olsen 1981), which suggests that the location of the retinal specialization may not overlap with the functional focus.

### Functional foci do not overlap with highest ganglion cell density

To determine whether the identified functional focus spatially overlaps with the area of highest density of retinal ganglion cells we made a mouse eye-model (Figure 3A), modified from previous models (Barathi, Boopathi et al. 2008). Using the eye model, retinal spatial locations could be projected through the optics of the mouse eye to the corneal surface. We first reconstructed the isodensity contours of published RGCs (Drager and Olsen 1981) to define the retinal location of the mouse retinal specialization (Figure 3- figure supplement 1A-C, note that these contours are also in agreement with other recently published maps of total RGC density (Zhang, Kim et al. 2012, Bleckert, Schwartz et al. 2014)). The lens optical properties were based on a GRIN lens (present in both rats (Philipson 1969, Hughes 1979) and mice (Chakraborty, Lacy et al. 2014)). To determine the optical characteristics of this lens we developed a method which combined models of the lens surface and refractive index gradient (Figure 3A, Figure 3- figure supplement 1D and Tables 1 and 2, see methods for details). Using this model, the contours representing the retinal specializations were projected through the eye model onto the corneal surface to determine equivalent corneal locations (Figure 3B, Figure 3- figure supplement 1E). Comparing this corneal projected location to the functional focus location showed that the region with the highest RGC counts had no overlap with the functional focus (Figure 3B) and occupied non-overlapping peripheral locations (Figure 3C). Viewed from above the animal’s head the functional foci were directed at the region in front of the animal’s nose and within the region of stable binocular overlap (azimuth: 1.4±8.8° and −4.4±9.3°, elevation 5.7±2.1° and 4.9±1.4° for left and right eyes respectively, N = 13641 frames, N=3 mice), while the retinal specialization was directed laterally (azimuth: −66.2 ±6.7° and 70.3±4.7°, elevation: 30.8±12.2° and 41.0±13.5° for left and right eyes respectively, N = 13641 frames N=3 mice. Figure 3D, Figure 3 – figure supplement 1F-G). As the projected location of the RGC high density region and the functional focus are both sensitive to torsional offsets and the location of the RGC region is also sensitive to the selected location for the optical axis of the eye model, we next measured what rotational transformations would be required for the RGC high density region and the functional focus to overlap. The size and locations of the two regions are such that there is no torsional rotation which would produce overlap (Figure 3 – figure supplement 1H-K). Any overlap of the regions would then require a large offset in the placement of the optical axis of the eye model on the redrawn retinal whole mount. Here we aligned the optical axis of the eye model with the center of mass of the redrawn optic disc, which has been measured as being 3.7° from the geometrical center of the retina (Sterratt, Lyngholm et al. 2013). As the spherical distance between the centers of the two regions was 52° (mean±SD, Left, 52.9±1.4°; Right, 51.4±4.6°, n=3 mice), no reasonable offsets or errors could result in overlap. Together this shows that that although mice maintain their prey within a focal visual region during the tracking phase of their pursuit behavior, this region does not overlap with the visual space represented by overall highest density RGC region of the retina (Drager and Olsen 1981, Jeon, Strettoi et al. 1998, Zhang, Kim et al. 2012). As a high-density of Alpha-ON sustained RGC’s are spatially located on the dorso-temporal retina (Bleckert, Schwartz et al. 2014), consistent with projecting to the front of the animal, we next quantified whether this region overlapped with the functional focus observed here (Figure 3E).

**Figure 3 with 1 supplement.**
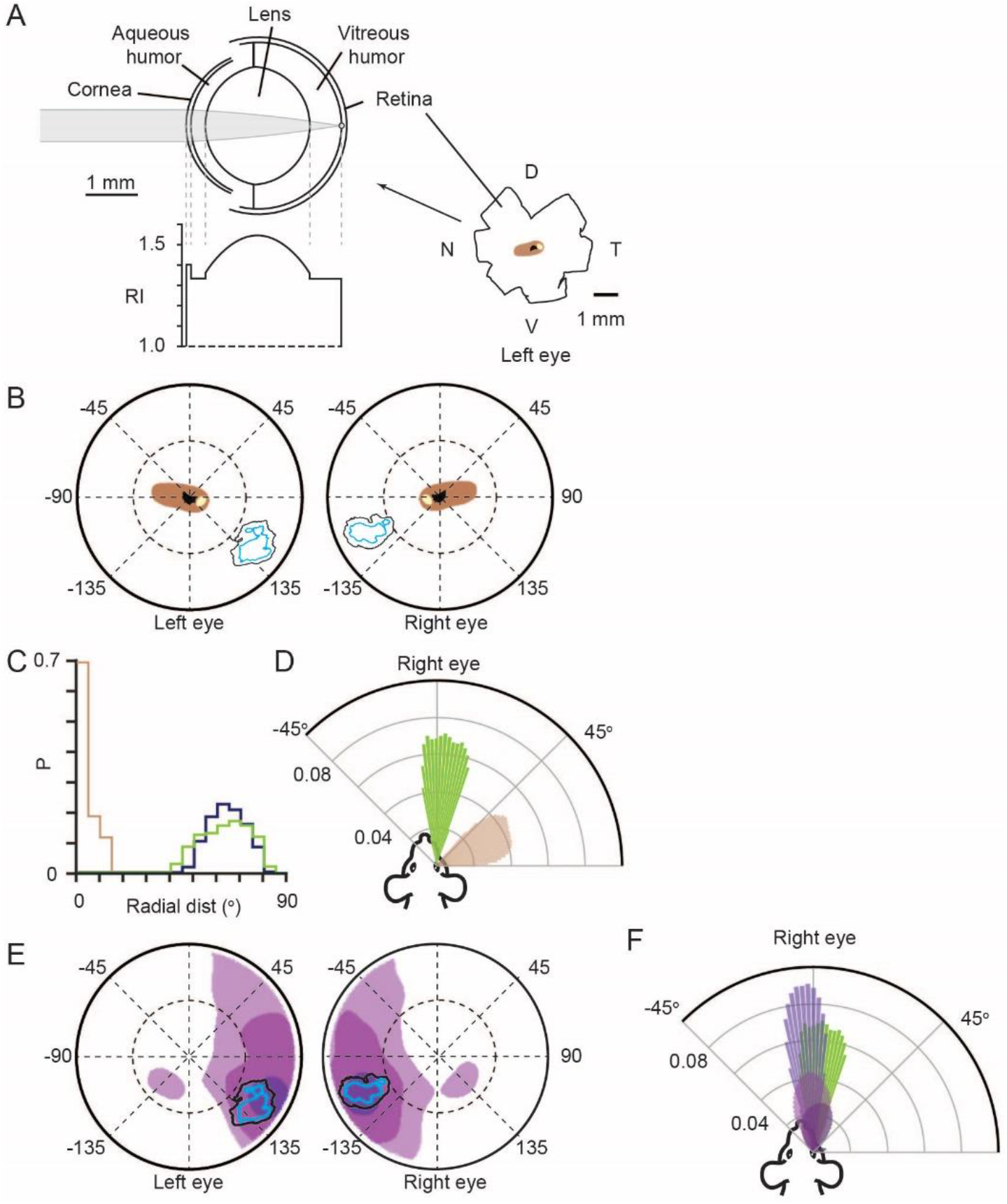
Functional foci are not sampled by the highest density retinal ganglion cell region. **(A)** Schematic of mouse eye model (left upper) with profile of all refractive indices (RI, left lower). Reconstructions of the optic disc (black), highest (>8000 cells/mm^2^, beige) and second highest (>7000 cells/mm^2^, brown) retinal ganglion cell (RGC) density regions redrawn from Drager & Olsen 1981, shown in lower right. **(B)** Position in corneal views of the high RGC density regions (brown and beige filled regions), and isodensity contours from Figure 2H after projection through the eye model. Orientation as in Figure 2E. **(C)** Horizontal axis histograms for the nasal half of the corneal view of the second highest RGC region (brown) and 50% isodensity contour for left (blue) and right (green) eyes. **(D)** Top-down view of the coverage regions for the right eye of the 50% isodensity contour (green, N = 7551 frames) and second highest RGC region (brown, N = 51007 frames) for a single animal. Bars represent the probability density function for the respective regions at that azimuth angle. **(E)** Position in corneal views of Alpha-ON sustained RGC densities (redrawn from Bleckert et al. 2014) after projection through the eye model. Colored regions show the 95% (dark purple), 75% (medium purple) and 50% (light purple) contour regions of the peak Alpha-ON sustained RGC density. lsodensity contours from Figure 2H. **(F)** Top-down view of the coverage regions for the right eye of the 95% (dark purple), 75% (medium purple) and 50% (light purple) Alpha-ON sustained RGC contour regions (same as in E, N = 51007 frames) and the 50% isodensity contour from D (green) for a single animal. For the Alpha-ON sustained RGC contour regions 50% means that this region contains all points which are at least 50% of the peak RGC density.

**Table 1.** Mouse eye model curvatures. Radii of curvature of the optical components of the mouse eye model in Figure 3A.

| *Ocular Component* | *Radius of curvature*<br>*($\mu\text{m}$)* |
| --- | --- |
| *Anterior Cornea* | -1408* |
| *Posterior Cornea* | -1372* |
| *Anterior Lens* | -1150* |
| *Posterior Lens* | 1134* |
| *Retina* | 1598* |

\* -values from (Barathi, Boopathi et al. 2008)

**Table 2.** Mouse eye model thicknesses and refractive indices. Parameters for the mouse eye model in Figure 3A.

| *Ocular Component* | *Thickness($\mu\text{m}$)* | *Index of refraction* |
| --- | --- | --- |
| *Cornea* | 92* | 1.402* |
| *Anterior chamber* | 278* | 1.334* |
| *Lens* | 2004* | 1.36 – 1.55<sup>#</sup> |
| *Vitreous chamber* | 609* | 1.333* |

\* - values from (Barathi, Boopathi et al. 2008)
<sup>#</sup> - minimum and maximum values after eye model optimization.

The average 50% contour of the functional focus was overlapped by the highest density of On Alpha-ON sustained RGC’s by 35 and 67% for left and right eye respectively (Figure 3E, black, mean±SD for left and right eye, 35.1±19.8 %, 66.7±0.09 %, p= 0.095 & 0.019, one-sided Student’s t-test), and for the overlap with the second highest density was 83 and 95% (mean±SD for left and right eye, 82.8±20.1 %, 94.8±24.7 %, p= 0.042 & 0.003, one-sided Student’s t-test), suggesting a high degree of correspondence between the highest density of Alpha-ON sustained RGC’s and the functional focus during pursuit behavior. Viewed from above the animal’s head the functional foci were directed at the region in front of the animal’s nose azimuth: 1.4±8.8° and −4.4±9.3°, elevation: 5.7±2.1° and 4.9±1.4° for left and right eyes respectively, N = 13641 frames, N=3 mice). The Alpha-ON sustained RGC’s were also directed in front of the animal’s nose (mean±SD, elevation:16.0±6.9° and 10.8±11.0 °, azimuth: −3.6±0.7 ° and 5.8±7.9 ° for left and right eyes respectively, N = 168400 frames, N=3 mice, Figure 3F). Together this suggests that objects falling within the functional foci are processed by Alpha-ON sustained RGC’s.

### Combination of torsional, horizontal, and vertical eye rotations counter head rotations

Eye movements in freely moving mice, like with rats (Wallace, Greenberg et al. 2013), can be large and rapid (Payne and Raymond 2017, Meyer, O’Keefe et al. 2020), and counter head rotations through the VOR (Figure 4 – figure supplement 1), enabling the large field of view around the animals head to be stabilized while the animal is moving. While the relationships between head rotations and both the horizontal and vertical eye rotations have been quantified, how these rotations combine with torsional rotations is not known. If mouse VOR operates similar to that observed in the rat (Wallace, Greenberg et al. 2013), torsional rotations in the mouse will play a significant role in stabilizing the visual field particularly during changes in head pitch. As with the vertical and horizontal rotations (Meyer, Poort et al. 2018), torsional rotations in freely moving mice spanned a wide range of rotation angles (Figure 4 – figure supplement 2A-D), and were correlated with head pitch (Pearson’s correlation coefficient (r): detect −0.72, 0.58, track: −0.60 and 0.53 for left and right eyes respectively, N=4406 detect and 13624 track frames, N=3 mice, Figure 4 – figure supplement 2C-D) as well as head roll (Pearson’s correlation coefficient (r): detect: −0.46, −0.47 track: −0.45 and −0.48 for left and right eyes respectively, N=4406 detect and 13624 track frames, N=3 mice, Figure 4 – figure supplement 2 L-M), as found previously for freely moving rats (Wallace, Greenberg et al. 2013). As with rats, the rotational relationship between the two eyes was dynamic with different forms of coordination (Figure 4 – figure supplement 2E-I), including episodes of in- and excyclovergence (torsional rotation of both eyes toward or away from the nose respectively) as well as dextro- and levocycloversion (torsional rotation of both eyes to the animal’s right or left respectively). We next analyzed how effectively rotations of the eye around multiple rotational axes combined to compensate the rotation of the head (Figure 4A, Figure 4 – figure supplement 3A-G). We compared movement of the head around its rotational axes and eye movements around the same rotational axes (Figure 4A), effectively defining alternative rotational axes for the eyes to match the axes of the head. Rotation of the eye around these re-defined axes would involve simultaneous rotations in multiple of the eye’s anatomical axes. The gain of this compensation was relatively linear and less than unity for both pitch- and roll-axes, indicating on average under-compensation of the head rotation (slope (gain) of relation for pitch axis, −0.45±0.12 and −0.48±0.06; roll axis −0.51±0.12 and −0.62±0.05 for left and right eye respectively, 168852 frames, N=3 mice). The relatively linear relationships between head and eye rotation around the head pitch and roll axes (Figure 4B) with a transition through the origin suggests that the horizontal, vertical and torsional eye movements are combined to effectively compensate pitch- and roll-related head movements. We next digitally froze each individual eye rotation axis (torsion, vertical and horizontal) and measured the effect on countering the head rotation (Figure 4C). For rotations around the head x-axis (head pitch changes) the gain of compensation was most affected by freezing torsional rotations (Figure 4C, gain mean±SD, control: −0.45±0.12and-0.48±0.06; torsion frozen −0.24±0.1 and-0.24±0.01, for left and right eyes respectively, N= 168852 frames, N=3 mice), while freezing vertical or horizontal rotations had more minor effects (Figure 4C, Table 3). The gain of compensation for rotations around the head y-axis (head roll changes) was dramatically affected by freezing vertical rotations (Figure 4C, gain mean±SD, control: −0.51±0.12and −0.62±0.05, vertical frozen −0.16±0.14and −0.17±0.03, for left and right eyes respectively, N= 168852 frames, N=3 mice), with freezing torsion also reducing compensation gain but to a lesser extent (Figure 4C, Table 3). We next quantified the stability and alignment of the animal’s binocular visual field during the pursuit sequences and determined the location of the functional foci within the stabilized region.

**Figure 4 with 3 supplements.**
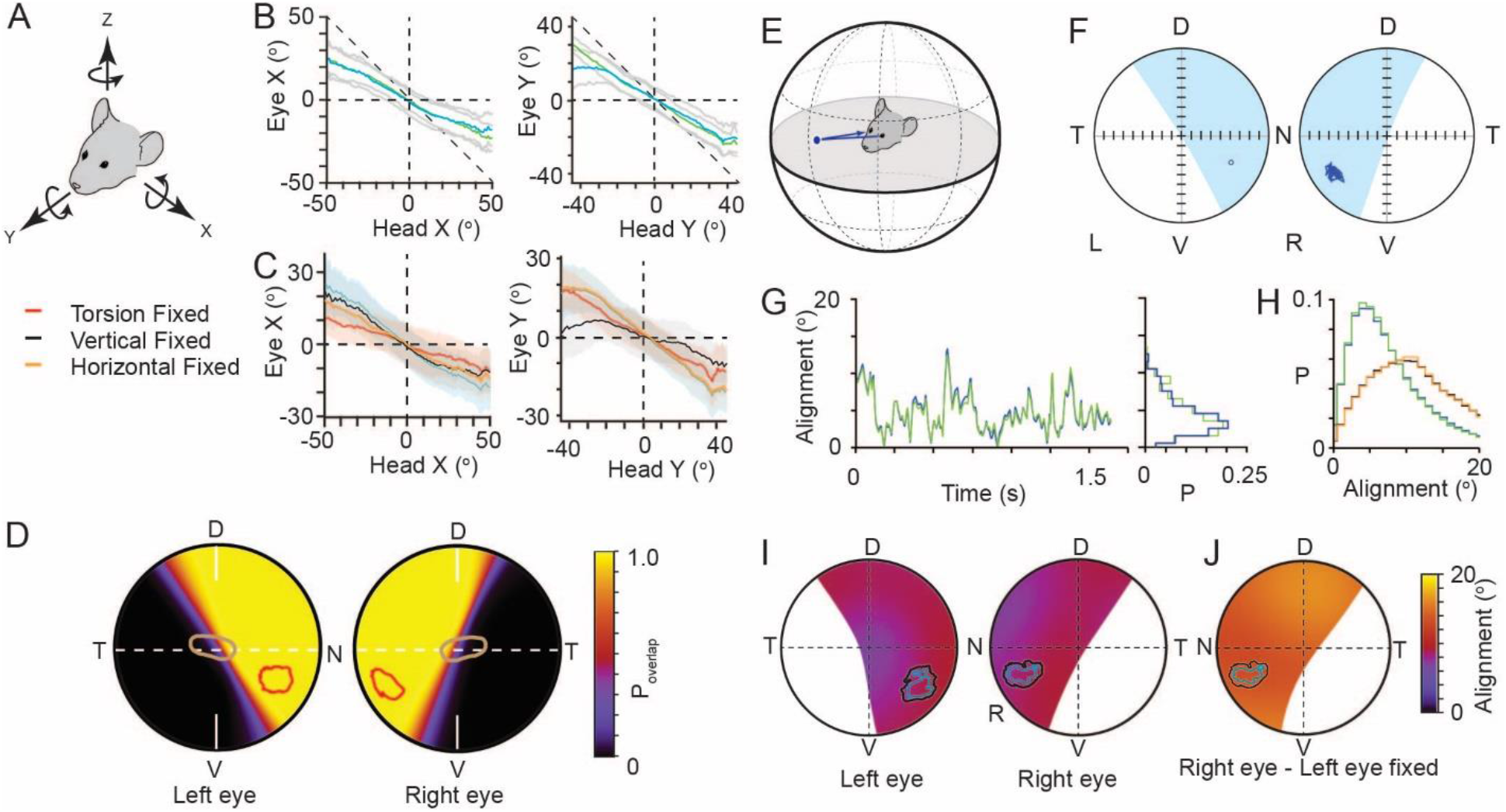
Functional foci are located within binocular regions in which motion is stabilized. **(A)** Schematic of the common head and eye rotational axes. **(B)** Relationship between head and eye rotations around the common X (left, 154500 frames from 3 animals) and Y (right, 165345 frames from 3 animals) rotational axes during pursuit and non-pursuit sequences. Plots show mean for left (blue) and right (green) eyes with standard deviation (gray). **(C)** Relationship between head and left eye rotations around the common X (left) and Y (right) rotational axes with; all eye rotations present (blue), torsional eye rotations frozen (red), vertical eye rotations frozen (black) or horizontal eye rotations frozen (orange). Plots show mean (lines) and standard deviations (colored filled regions). **(D)** Corneal view showing probability of overlap of left and right visual fields for one example animal (71995 frames), with overlay of isodensity contours (red) from functional foci (see Figure 2 - figure supplement 1D) and contour of second highest RGC region (brown) from Figure 3B. **(E)** Schematic of inter-ocular alignment. **(F)** Corneal view of alignment reference point in left eye (left) and variability in alignment of the re-projection of that point in the right eye (right) for a 1.6s data segment. **(G)** Kinetics (left) and associated distribution (right) of the variability in ocular alignment for left eye point projected to right eye (blue) and right eye point in left eye (green) for one example data segment (shown in G) from one animal. **(H)** Distributions of ocular alignment from all data segments (159318 frames, n==3 mice) with the measured eye movements for left into right eye (blue) and right into left eye (green) and alignment with eye movements frozen (left into right eye, black, right into left, orange). **(I)** Map of average inter-ocular alignment for all data segments (159318 frames, n==3 animals) with overlay of isodensity contoursfrom Figure 2H. **(J)** Map of average inter-ocular alignment as in J with left eye movements frozen.

**Table 3.** Compensation gain of eye rotations for head X or Y-axis rotations. Effect of digitally freezing torsional, vertical and horizontal eye rotations on the gain of compensation of X and Y head rotations. Data taken from 168852 frames, from 3 animals

| Eye | Rotation direction | Rotation | All Rotations<br>(mean ± SD) | Eye rotation frozen<br>(mean ± SD) |
| --- | --- | --- | --- | --- |
| Left | X | Torsion | -0.45±0.12 | -0.24±0.1 |
| | | Horizontal | -0.45±0.12 | -0.32±0.06 |
| | | Vertical | -0.45±0.12 | -0.35 ±0.08 |
| Right | X | Torsion | -0.48±0.06 | -0.24±0.01 |
| | | Horizontal | -0.48±0.06 | -0.36±0.08 |
| | | Vertical | -0.48±0.06 | -0.34±0.03 |
| Left | Y | Torsion | -0.51±0.12 | -0.35±0.05 |
| | | Horizontal | -0.51±0.12 | -0.51±0.11 |
| | | Vertical | -0.51±0.12 | -0.16±0.14 |
| Right | Y | Torsion | -0.62±0.05 | -0.45±0.05 |
| | | Horizontal | -0.62±0.05 | -0.62±0.02 |
| | | Vertical | -0.62±0.05 | -0.17±0.03 |

### Functional foci are located in the motion-stabilized binocular visual field

Similar to rats, left and right visual fields overlapped extensively (Hughes 1979, Drager and Olsen 1980), with eye movements creating variability in the extent of the overlap at the edges of the two visual fields, the transition from monocular to binocular (Figure 4D). The functional foci for both eyes were predominately contained within the region of continuous binocular overlap. A horizontal transect through the optical axis for all animals showed a gradual transition from continuous binocular coverage to zero binocular coverage commencing just nasal of the optical axis (Figure 4D, Figure 4 - figure supplement 3H and I), indicating that the region of highest RGC density spans this transitional region whereas the functional foci are, on average, contained within the binocular region (Figure 4D - figure supplement 3H).

We next quantified the variability of alignment of the left and right visual fields within the binocular region, and specifically in the functional focus location (Figure 4E) by using the center of mass (50% isodensity contour center) of the left eye functional focus as an initial reference point and projecting this point to the boundary of a hypothetical sphere surrounding the head. This contact point on the sphere was then re-projected into the right eye to identify the matching location of the left eye (Figure 4E). We then followed the trajectory of the re-projected point in the right eye to get a measurement of alignment variability (Figure 4F, for comparison with the locations in the right eye projected into the left eye see Figure 4 – figure supplement 3I-K). While pursuing crickets, alignment precision varied through time (Figure 4G) with the mean alignment of the reference point over all animals and data segments being ~8-9°, which is around the size of V1 cortical neuron receptive fields (~5-15° (Niell and Stryker 2008), Figure 4H, mean±SD, left eye projected into right eye 8.8±6.9°; right eye projected into left eye 8.6±6.7°). Repeating this analysis for all points within the region where the probability of binocular overlap was greater than 5% showed that there was a relatively uniform alignment over the entire region (Figure 4I), and that the average alignment error in the functional foci was 8-10°. Coordination of eye movements was important for alignment, as freezing the movements of one eye to its mean position resulted in a significant increase in the alignment error when comparing individual cricket tracking sequences (left all rotations vs. left eye frozen P=1.78x-10, right eye all rotations vs. right eye frozen P=7.12×10-11, N=52 sequences, unpaired Student’s t-test), and a ~54% increase in the mean alignment error over all frames for the reference location (Figure 4I, left eye projected into right eye (left eye frozen) 13.4±8.3°; Right eye projected into left eye (right eye frozen) 13.4±8.3°, mean±SD, 159318 frames, N=3 mice), which also resulted in a uniform increase in alignment error over the whole overlap region (Figure 4J and Figure 4 – figure supplement 3J-L). In summary, during pursuit behavior the functional foci are located in a stable binocular region of the mouse’s visual field. However, in the absence of a mechanism for voluntarily directing its gaze towards a specific target, such as smooth pursuit, tight coupling of VOR evoked eye movements to head rotations would seem to be restrictive to an animal’s ability to move the target into a specific part of their visual field during pursuit. We therefore next measured what mechanisms mice use to bring the prey into their functional focus.

### Behavioral mechanisms for maintaining prey within functional foci

At detection, mice orient towards their target, aligning their head with the prey and running towards it (Figure 2D), keeping the cricket within a narrow window around its forward direction. This provides a direct way for mice to hold their prey within their binocular visual fields (Figure 4D). We next measured whether additional head or eye movements are used to keep the target within the functional foci. If the mice were actively maintaining the prey within a fixed location of their visual fields the position of the cricket image should not change as the mouse approaches the cricket. The cricket image location could be maintained by either a head or eye rotation. If they were not actively maintaining the prey in a fixed location, the cricket images should shift downwards in the visual fields as the mouse approaches. To distinguish between these two possibilities we plotted the cricket positions in the eye views color-coded by the distance between the mouse and cricket (Figure 5A). As the mouse approached the cricket during the track behavioral epoch, the projected cricket positions shifted lower in the visual field (Figure 5A lower). This suggests that the mice did not use additional head or eye movements (Figure 5 – figure supplement 1) to bring the cricket into the functional foci, but rather manipulated the cricket’s position in the eye view by orienting and moving towards the target. Consistent with this, head pitch remained stable as the mice approached the crickets (Figure 5B). Furthermore, there was no significant difference in head pitch as a function of distance to the cricket between non-tracking and tracking periods (non-tracking head pitch: −3.7±26.5°, mean±SD, median = −11.3 °, tracking head pitch: −12.9±15.7°, mean ± SD, median = −14.6°, Ks test, P=0.709, paired Student’s t-test P=0.197, N=18 detect-track sequences, N= 3 mice). In addition, and consistent with previous findings (Michaiel, Abe et al. 2020), mice did not significantly converge their eyes as they approached the crickets (non-tracking head vergence: 7.6 ±13.5°, mean±SD, median = 8.6°, tracking head vergence: 2.5 ±16.7°, mean ± SD, median = 3.2°. Ks test, P=0.425, paired Student’s t-test P=0.225, N=18 detect-track sequences N = 3 mice, Figure 5 – figure supplement 1J and Table 4). These observations suggest that the primary role for the eye movements is stabilizing the visual fields.

**Figure 5 with 1 supplement.**
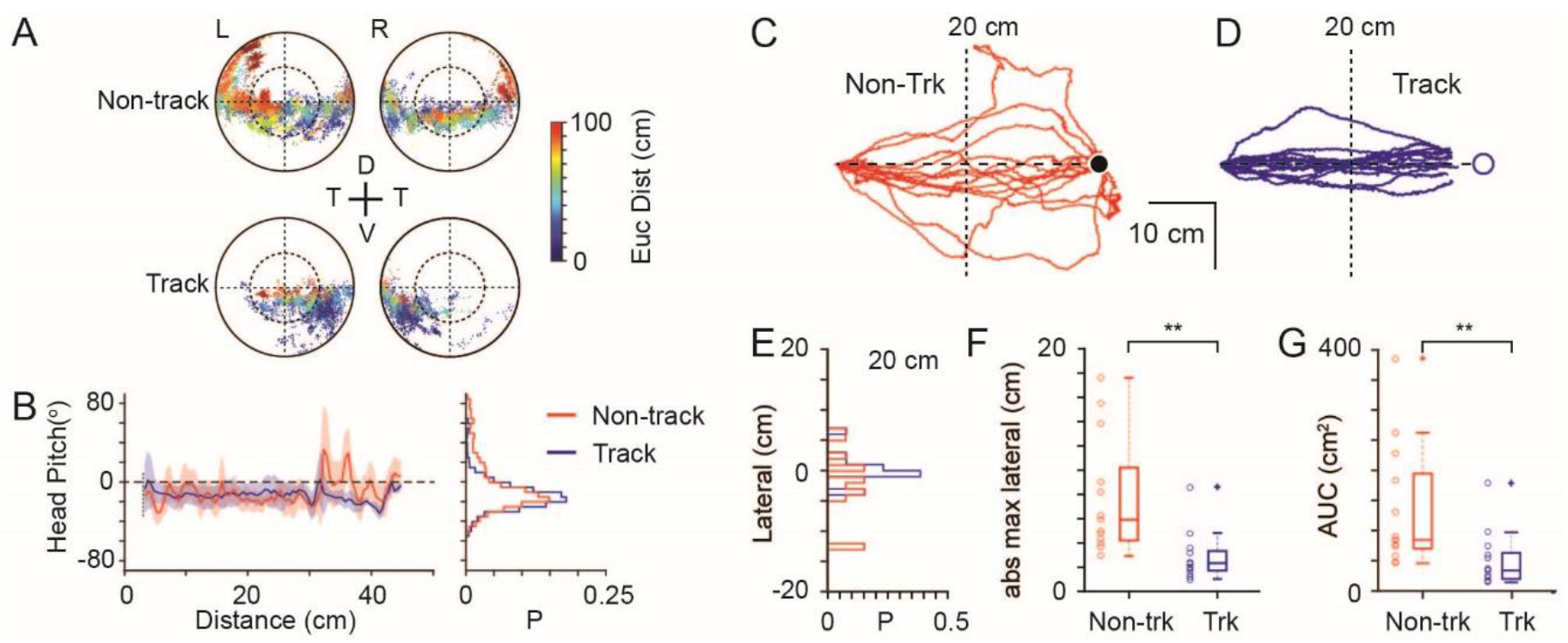
Mechanisms used to maintain prey within a focal visual region. **(A)** Corneal locations of cricket position color-coded by Euclidean distance to cricket for non-track (upper) and track (lower) epochs (18 data sequences, 15649 non-tracking and 8510 tracking frames, n=3 animals). **(B)** Mean and SD head pitch with Euclidean distance to cricket (left) and distribution of head pitch angles (right) for non-track (red) and track (blue) epochs (datasets as in A). **(C)** Mouse trajectories during non-track epochs rotated and overlaid to show deviation from a direct path (13 trajectories from 3 animals). **(D)** Mouse trajectories as in D but during track epochs (13 trajectories from 3 animals). **(E)** Histogram of lateral deviations for non-track (red) and track (blue) data in C and D calculated 20 cm from the end of the trajectory. **(F)** Boxplots and individual data points of absolute maximal lateral deviation from a direct path between start and end points for non-track (red) and track (blue) epochs (datasets as in C & D), ** P=0.0006, Wilcoxon’s Rank Sum Test. **(G)** Boxplots and individual data points of area under the curve (AUC) of mouse trajectories during non-track (red) and track (blue) epochs (datasets as in C & D),** P=0.0029, Wilcoxon’s Rank Sum Test.

**Table 4.** Eye rotations during non-tracking and tracking periods. Horizontal, vertical and torsional eye rotations during the non-tracking and tracking periods in Figure 5. Data taken from 18 non-track epochs and 18 track epochs, from 3 animals.

| Ocular Rotation | Non-Trk<br>(mean ± SD)<br>(median) | Track<br>(mean ± SD)<br>(median) | p value (KS) | P value<br>(Student<br>T-test) |
| --- | --- | --- | --- | --- |
| Lt Horizontal | -1.8 ± 9.9°<br>(-1.7°) | -1.8 ± 14.9°<br>(-3.5°) | 3.9x10<sup>-2</sup> | 0.162 |
| Lt Vertical | 0.8 ± 11.2°<br>(-0.4°) | 4.5 ± 11.1°<br>(4.9°) | 0.425 | 0.616 |
| Lt Torsional | 2.9±16.1°<br>(0.0°) | 1.3±20.6°<br>(0.0°) | 0.945 | 0.610 |
| Rt Horizontal | 5.7 ± 10.9°<br>(5.5°) | 1.0 ± 9.9°<br>(1.7°) | 9.82x10<sup>-2</sup> | 1.08x10<sup>-2</sup> |
| Rt Vertical | -3.6±13.4°<br>(-6.3°) | 5.6±12.7°<br>(-7.1°) | 0.945 | 0.804 |
| Rt Torsional | 0.32±13.5°<br>(0.0°) | 0.7±12.3°<br>(0.0°) | 0.425 | 0.366 |

If mice successfully track and capture prey by retaining the target in front of them, then this should be reflected in the trajectories taken by the mice during the tracking epoch compared to the non-tracking behavioral epochs. During cricket tracking periods, mice ran directly towards the cricket, and their trajectories were significantly straighter than during equivalent non-tracking phases (Figures 5C-G). Lateral deviation at the half-way point (Figure 5E, non-tracking 4.3±4.0 cm, tracking 1.4±2.0 cm, P=0.009), maximum lateral deviation (Figure 5F, non-tracking, 7.7±4.9 cm, tracking 2.8 ±2.0 cm, P=0.0006) and the area between the trajectory and minimum distance path to the target (Figure 5G, area under the curve, non-tracking 135.6±102.7 cm^2^, tracking 51.3±45.8 cm^2^, P=0.0029) were all significantly smaller in the tracking epochs (all comparisons mean±SD, N=13 tracking and non-tracking sequences, N=3 mice, Wilcoxon’s Rank Sum Test).

Together this suggests that mice do not make compensatory vertical head movements, tracking eye movements or vergence eye movements to keep prey within their functional foci but instead retain their target within a restricted bearing by running straight towards it. This raised the question of what advantage is this behavior to the mouse and what is unique about the functional focus position on the cornea?

### Functional foci are located in region of minimized optic flow

Optic flow is the pattern of object motion across the retina that can be self-induced, through eye, head or translational motion, or induced by motion of objects in the environment, or combinations thereof (for review see: (Angelaki and Hess 2005). In a freely moving animal in a still environment, translational motion results in a pattern of optic flow that consists of a radial flow-fields emanating from a point of zero-velocity (Figure 6A). While optic flow is used by many species for both navigation and the estimation of the motion properties of moving objects, motion induced blur degrades image formation on the retina and decreases resolution depending on the animal’s direction of travel (Land 1999). Optic flow is minimized in the direction of travel directly in front of the animal (Sabbah, Gemmer et al. 2017), with flow fields directed away from travel direction and forming a second minimum directly behind the animal’s head (Figure 6A, see also (Angelaki and Hess 2005). To measure the characteristics of optic flow in in both eyes of freely moving mice and to relate this flow pattern to the location of the functional foci, we next calculated average optic flow from freely moving data during pursuit behavior using the digitized environment and eye-views (Figure 6B). First, we calculated optic flow in the idealized case of forward translation motion when all surrounding surfaces were equidistant (Figure 6C). As mice have laterally facing eyes (optical axis = 59.9±19.8° and −62.3±14.7° lateral of frontal for the left and right eyes respectively, N=3 mice), idealized forward motion resulted in the region of minimal optic flow in each eye being located off optical axis in the ventro-medial corneal region representing the animal’s forward direction (radial displacement from optical axis 36.64±0.92° and −41.11±2.27°, rotational angle 122.95±17.05° and −107.94±9.96°, for the left and right eyes respectively, mean ± SD, N=2 mice, Figure 6C). During free movement both the distance from the eyes to objects in the environment, as well as head and eye-rotations had a strong influence on the optic flow fields. We visualized the average flow fields during free motion by calculating the optic flow on the cornea during multiple pursuit trials (N=20 prey chases containing 52 tracking sequences, initial Euclidean distance mouse-cricket >20 cm). The resulting optic flow density maps were complex with a wide range of average speeds (133.44±221.42 °/s, mean±1SD, median 28.64 °/s, interquartile range 4.57–137.18 °/s, N=2 mice, Figure 6D). The area of lowest optic flow extended from nasal field of view to overhead (Figure 6D) but unlike the simulated case (Figure 6C) optic flow was not symmetric around the regions of minimal optic flow. Optic flow in the 30×30° region surrounding the ventro-medial point of minimal optic flow was significantly lower than that in an equivalent region in the ventro-temporal region during free movement, but not in the simulated case (free movement: nasal 46.3±9.8 °/s, temporal: 199.4±29.0 °/s, P=0.0014, simulated: nasal 163.6±82.2 °/s, temporal: 833.0±416.5 °/s, P=0.0662, mean±SD, two-sided t-test, unequal variance, N=2 mice). Optic flow was higher in the lower visual field and considerably lower in the upper visual field (lower left eye visual field: 262.44±106.50 °/s, upper left eye visual field: 44.87±24.31 °/s, P=1.78×10^−20^, lower right eye visual field: 361.91±168.80 °/s, upper right eye visual field: 40.59±22.79 °/s, P=6.68×10^−19^, Two-sided t-test, unequal variance, N=2 mice), due to the greater distance between ceiling and mouse (distance to floor 2±1cm, distance to ceiling 308±107cm, 9873 frames, N=3 mice). Given the advantage of low optic flow to mammalian vision, we next quantified the position of least optic flow during prey tracking. We calculated the location of the translational optic flow minimum in each frame for each eye, and created a probability map of this location over the visual field (Figure 6E). The region of highest likelihood for the presence of the optic flow minimum overlapped considerably with the functional foci in both eyes during the tracking epochs of the pursuit behavior (overlap of optic flow 95% minima and functional foci 50% regions: 100% and 99±1%, overlap of optic flow 50% minima and functional foci 50 % regions: 61±14 % and 72±4 % in left and right eyes respectively, N=3 mice, Figure 6E). Together this shows that mice preferentially maintain their prey in the region of reduced optic flow during pursuit, where the retinal image of their prey is least distorted due to motion induced image blur.

**Figure 6.**
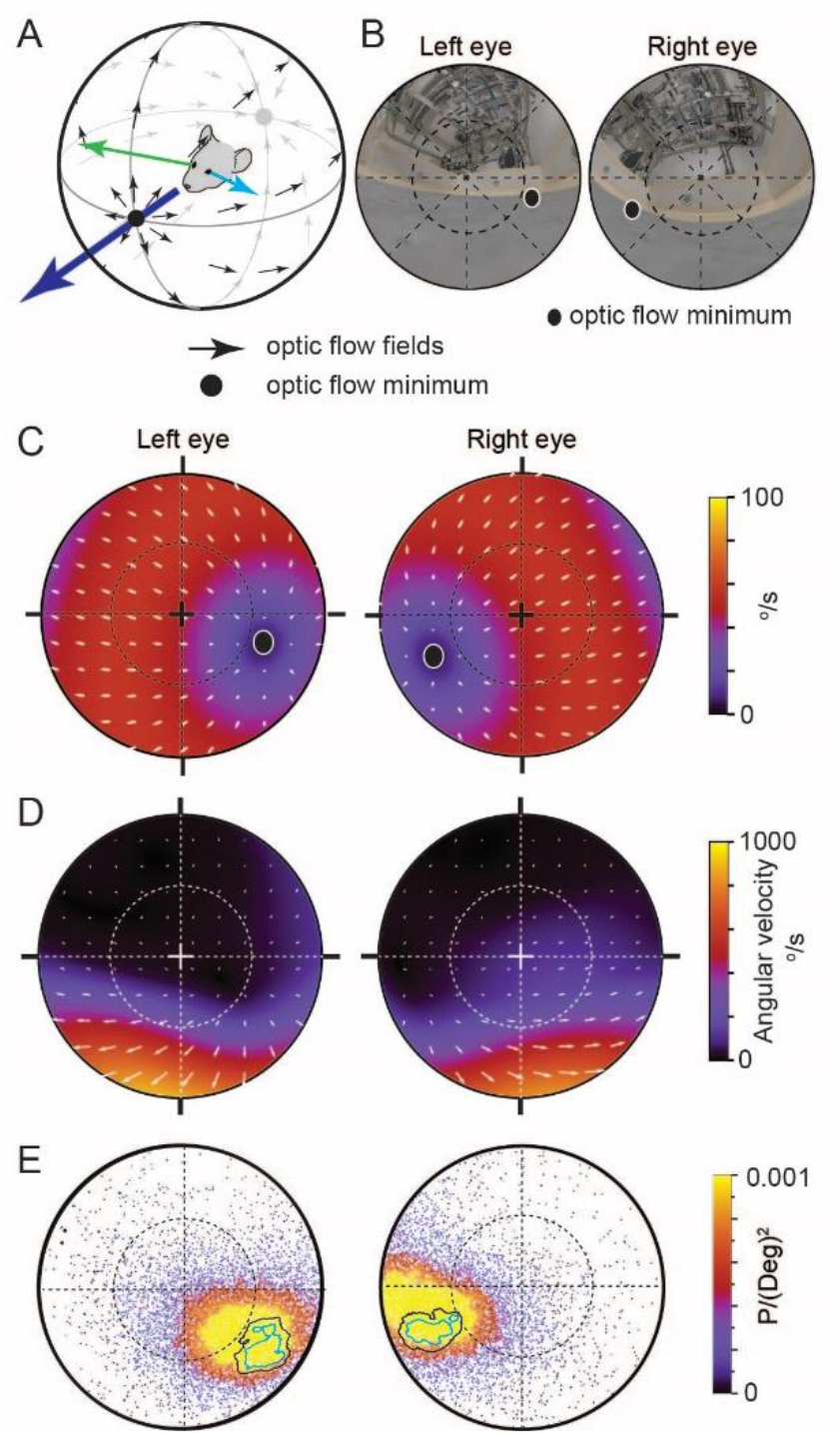
Functional foci are located in the regions of reduced optic flow during forward motion. **(A)** Schematic of idealized optic flow (black arrows) as a mouse translates forwards (after (Sabbah, Gemmer et al. 2017)). Left (blue arrow) and right (green arrow) gaze vectors. **(B)** Location of optic flow minima in reconstructed mouse eye views of the cricket and experiment arena (from Figure 1H), circle represents 45°. **(C)** Optic flow map in corneal views, showing flow velocity (color coding) and direction (white arrows) calculated for the idealized spherical environment in 6A with forward motion of 50 cm/s. **(D)** Optic flow maps in corneal views during track epochs (5269 frames), from one animal. **(E)** Probability density map of optic flow poles in mouse corneal views during track epochs (data as in Figure 2G, 13641 frames), with overlay of isodensity contours from Figure 2H.

## Discussion

We developed a technique for reconstructing the visual fields in a freely moving mouse during prey pursuit to quantify the spatial relationship between the environment, cricket and the mouse. Using this approach, we show that mice, while pursuing crickets, preferentially maintain the prey in a localized region of their visual field, termed here the functional focus. The positional maintenance of the cricket was not achieved by active eye movements that followed the prey, but rather by the animal’s change in behavior, specifically the head-movement and orientation towards the prey during pursuit. While eye rotations stabilized the visual field via the vestibulo-ocular reflex by countering head rotations, the rotations were not specific to either prey detection or prey tracking. This strongly suggested that eye-rotations in mice, like in rats, primarily stabilize their large field of view and that all three rotational axes, including ocular torsion, combine to counter head rotations. In addition, we also show that eye rotations cannot be predicted from head rotations in any one axis as has been suggested by recent studies of mouse eye motion (Meyer, Poort et al. 2018, Meyer, O’Keefe et al. 2020, Michaiel, Abe et al. 2020) but rather by a combination of all head rotations (Figure 4 – figure supplement 2). As the eye rotations were predominately associated with countering head-rotations, this raised the question of whether the mouse can use a large fraction of its stabilized visual field to pursuit crickets, or whether a specific region is utilized. To accurately determine the correspondence between the animal’s visual field and the retinal image, we developed a quantitative model of the mouse eye and optics. Using this, we show that the location of the functional focus does not coincide with the retinal region with the highest total density of retinal ganglion cells that are laterally facing, but rather the highest density of Alpha-ON sustained RGCs, whose general properties have been previously proposed to be well suited for this purpose (Bleckert, Schwartz et al. 2014). Finally, we used the detailed, digitally rendered reconstruction of the arena and surrounding room to calculate the realistic optic flow in the visual fields (Gibson, Olum et al. 1955, Sabbah, Gemmer et al. 2017, Saleem 2020) of the mice as they pursued crickets, which showed that the functional foci coincide with the region of the visual fields with minimal optic flow during the cricket pursuit, and presumably are thereby minimally distorted by motion-induced image blur (for review see (Angelaki and Hess 2005)). Critical to this finding was the ability to isolate the visual sense, generate both a detailed reconstruction of both the local environment and the animal’s ocular anatomy and optical pathways, but also record eye motion in all three optical axes especially ocular torsion, something that has only been achieved in rats (Wallace, Greenberg et al. 2013). Lastly, by building an optical model and establishing the relationship between the retinal surface and the corneal surface we were able to relate the data generated from published studies on retinal anatomy (Drager and Olsen 1981, Sterratt, Lyngholm et al. 2013, Bleckert, Schwartz et al. 2014) and physiology (Pang, Gao et al. 2003, Murphy and Rieke 2006, van Wyk, Wassle et al. 2009, Dhande, Stafford et al. 2015, Martersteck, Hirokawa et al. 2017, Sabbah, Gemmer et al. 2017) to our behavioral data.

Both estimates of the field of view of the mouse eye (Drager 1978) and electrophysiological measurements of receptive field locations of visually responsive neurons (Drager and Olsen 1980, Wagor, Mangini et al. 1980) have established that the binocular region of the visual field in mice is contained within the nasal visual field of each eye, and spans a region of 30-40° in front of the animal (Wagor, Mangini et al. 1980). We present here, that similar to the rat (Wallace, Greenberg et al. 2013), the overlapping monocular fields that make up the binocular overlap are not constantly maintained (Figure 4H) but fluctuate at the margins as the eyes rotate to counter head rotations (Figure 4D), resulting in a region where there is a transition from one area with near continuous binocular coverage, through to a region that is invariably monocular. The functional focus described here lies within the region of high probability of maintained binocular overlap. This region of the visual field projects onto the temporal retina, which contains both ipsilaterally projecting (uncrossed) RGCs (Drager and Olsen 1980, Reese and Cowey 1986) and RGCs which form part of the callosal projection pathway (Olavarria and van Sluyters 1983, Laing, Turecek et al. 2015, Ramachandra, Pawlak et al. 2020), both of which are considered central to binocular visual processing. In addition, the current study adds to the significance to these previous findings and suggests that the functional focus location is well placed to support stereoscopic depth perception, assuming that this form of visual processing is available to and employed by the mouse (Scholl, Burge et al. 2013, Scholl, Pattadkal et al. 2015, La Chioma, Bonhoeffer et al. 2019, Samonds, Choi et al. 2019, La Chioma, Bonhoeffer et al. 2020). In addition, while the overall highest density of retinal ganglion cells in mice is located in the region around the optical axis (Drager and Olsen 1981), a recent study examining the distributions of various different subclasses of RGCs has shown that the highest density of Alpha-ON sustained RGCs resides in the superior-temporal retina (Bleckert, Schwartz et al. 2014) in a region which would approximately coincide with the functional focus. These Alpha-ON sustained RGCs have center-surround receptive fields, a rapid response and fast conducting axon, and are thought to be “spot detectors” (for review see (Dhande, Stafford et al. 2015)). In addition, the Alpha-ON sustained RGCs in this particular retinal region differ from the same RGC-type in other regions of the retina as they have a significantly smaller dendritic tree radius and subtend a smaller area of physical space as well as have overlapping receptive fields (Bleckert, Schwartz et al. 2014). Taken together, the cellular properties as well as the region in-front of the animal which provides their input are consistent with the requirements for tracking small and mobile targets (Lettvin, Maturana et al. 1959, Dean, Redgrave et al. 1989, Bleckert, Schwartz et al. 2014, Procacci, Allen et al. 2020). A recent study has shown that both wide-field and narrow-field neuronal types in the mouse superior colliculus play central roles in the detection and pursuit phases of this pursuit task respectively (Hoy, Bishop et al. 2019), and consistent with this, Alpha-ON sustained RGCs having projections to the superior colliculus (Martersteck, Hirokawa et al. 2017). It is currently unclear how the primary visual cortex (V1) contributes to this behavior, but some role is possible if not probable, which would also be supported by the strong Alpha RGC projection to the dorsal lateral geniculate nucleus and thus V1 (Martersteck, Hirokawa et al. 2017). Additionally, an increased cortical magnification factor occurs in the region corresponding to the nasal, binocular visual field (Schuett, Bonhoeffer et al. 2002, Garrett, Nauhaus et al. 2014).

Finally, we show that the region that contains these Alpha-ON sustained RGCs also coincides with the region of minimum optic flow and therefore reduced image blur during translation pursuit, a feature which would supports accurate localization of small targets by Alpha-ON sustained RGCs. Patterns of optic flow are thought to be an important component of perception of self-motion (Lappe, Bremmer et al. 1999). Mechanistically supporting this, global alignment across the retina of the preferred orientation of direction-selective retinal ganglion cells with the cardinal directions of optic flow during idealized motion has been shown in mice (Sabbah, Gemmer et al. 2017). The average optic flow measured here was, perhaps not surprisingly, strikingly different from that observed with idealized motion, resulting in large part from the large differences to objects in the environment in which the behaviors were performed. For fast moving, ground dwelling animals like mice, considerable asymmetry in optic flow across the visual field may be the more normal case, considering that objects above the animal are, in general, likely to be more distant.

In freely moving rats it has been shown that ocular torsion is correlated with head pitch such that nose-up rotation of the head is counteracted by incyclotorsion (rotation towards the nose) of both eyes, with nose-down pitch counteracted by excyclotorsion (Wallace, Greenberg et al. 2013). These rotations have the effect of stabilizing the horizontal plane of the retina with respect to the horizon. The considerable radial separation between the optical axis of the eye and both the functional foci observed in the current study as well as the highest density region of Alpha-ON sustained RGCs (Bleckert, Schwartz et al. 2014) renders the direction in which these regions point highly sensitive to torsional rotation. Consequently, torsional rotation also has an important effect on alignment of the left and right visual fields in addition to its role in visual field stabilization. We show here that torsional rotation in freely moving mice is also dynamic, with episodes showing in- and excyclovergence as well as dextro- and levocycloversion. Further, while the correlation between torsional rotation and head pitch observed in rats was measured, there was also an additional relation between ocular torsion and head roll consistent with VOR-evoked dextro- and levocycloversion. Consequently, prediction of torsion using a model based on head pitch alone resulted in an average error of around 7°, while an expanded model including roll as well performed better (Figure 4 – figure supplement 2J-O).

In summary, we show here that during pursuit mice preferentially keep their intended prey in a localized region of their visual fields, referred to here as the functional focus, but do so by orientating their head and body and running directly towards the prey rather than with specific eye movements. The location of the functional focus is within the binocular visual field, but in addition also coincides with the region of minimal optic flow during the pursuit, and presumably also minimally distorted by motion blur.

## Methods

### Animal details

Experiments were carried out in accordance with protocols approved by the local animal welfare authorities (Landesamt für Natur Umwelt und Verbraucherschutz, Nordrhein-Westfalen, Germany). Experiments were carried out using male C57/BL6JCrl mice (acquired from Charles River Laboratories). At the time of the cricket hunting experiments, mice (n=9) were between 2-8 months old, and weighed between 21-29g. Mice were maintained on a 12 hr light/dark cycle. Crickets (*Acheta domesticus*, Bugs-International, Germany) were housed in 480×375×210 cm cages with *ad lib* water and food (powdered mouse chow). Cricket body sizes ranged from 1 cm to 2 cm (1.8 ± 0.3 cm, mean ± SD, n=25).

### Implant surgery

Animals were anaesthetized using fentanyl, medetomidine and midazolam (respectively 50μg/kg, 5mg/kg and 0.5mg/kg, delivered i.p.), and analgesia was provided with carprofen (7mg/kg delivered s.c.). Body temperature was maintained using a thermostatically regulated heating pad. Respiration rate and depth of anesthesia was monitored throughout the procedure. Following opening of the skin and removal of connective tissue overlying the sagittal suture and parietal bones, the skull was cleaned with H_2_O_2_ (3%). A custom-made implant, consisting of a flat circular attachment surface for attachment to the skull, and implant body with three anti-rotation pins and a magnet (Figure 7A-B), was fixed to the dried skull using a UV-curing dental adhesive (Optibond FL, Kerr Corporation, Orange, California, USA) and a UV-curing dental composite (Charisma, Kulzer GmbH, Hanau, Germany). The implant attachment surface and body were made from light-weight, bio-compatible dental resin (Dental SG, Formlabs, Germany). Skin margins were closed with 5/0 Vicryl sutures (Ethicon Inc, Somerville, NJ, USA) and a cyanoacrylate adhesive (Histoacryl, B. Braun, Melsungen, Germany). The injectable anesthetic combination was antagonized with naloxone, atipamezole and flumazenil (respectively 1.2mg/kg, 0.5mg/kg and 0.75mg/kg, delivered i.p.), and the animal was allowed to recover.

**Figure 7.**
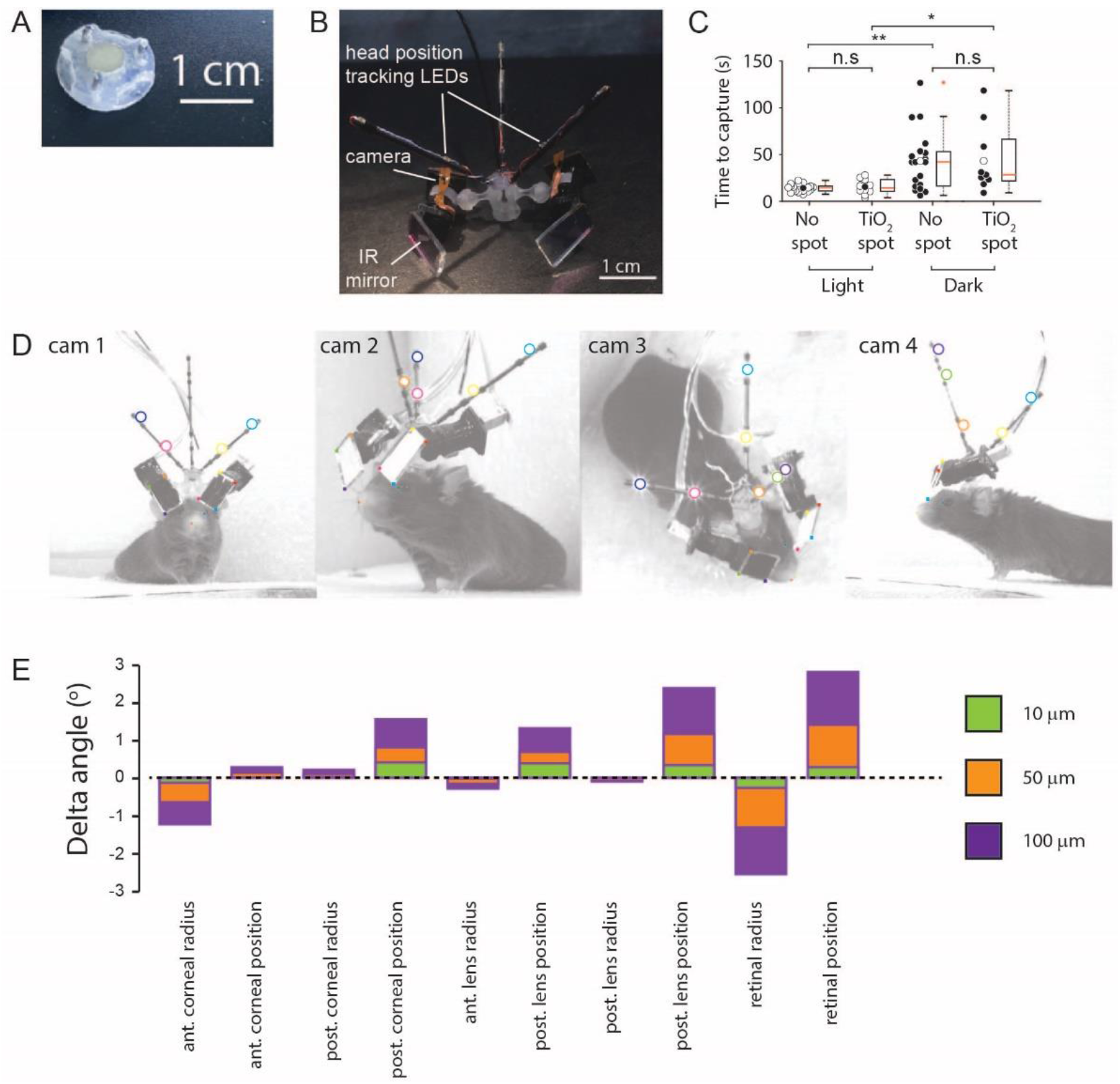
Methods. **(A)** Implanted baseplate with magnetic attachment point and restraining pin holes. **(B)** Miniaturized eye cameras and head position tracking system. Infrared illumination LEDs were mounted on the camera objective and reflected onto the eye using an IR-reflective mirror. Head position tracking IR-LEDs were mounted on three carbon-fiber struts attached to the head-mount. **(C)** Cricket capture times in lit or dark conditions in mice without (n=19 pursuit sequences, n=6 mice) or with (n=10 pursuit sequences in lit conditions and n=9 pursuit sequences in the dark, n=3 mice) corneal TiO2 torsion tracking spots, Lit vs Dark with no spot, P=0.0012, Lit vs Dark TiO2 spot, P=0.0133, Lit without spot vs Lit with TiO2 spot, P=0.69, Dark without spot vs Dark with TiO2 spot, P=1. n.s. = non-significant, *P<0.05, **P<0.01. Paired Wilcoxon’s signed rank tests. For these experiments, pursuits were conducted in a smaller arena (480 x 375 x 210 cm). **(D)** Images of mouse with eye camera and head position tracking system for anatomical calibration. Head mount and anatomical features marked. Anatomical features: Left (blue filled circles) and right (green filled circle) medial canthi, left (orange filled circles) and right (red filled circle) nostril positions. Head mount features: position tracking LEDs (large colored circles), IR mirror corner positions (small colored filled squares). **(E)** Sensitivity of the radial elevation on the retina in the mouse eye model to changes in the radii of curvature and thicknesses of the model optical components.

### Positioning of the head-mounted cameras

The eye cameras for oculo-videography were mounted on mounting arms which were attached to a baseplate with complementary holes to the anti-rotation pins on the implant and fitted with a magnet of complementary polarity. During positioning of the head-camera, mice were anaesthetized with isoflurane (induction: 3-5% isoflurane, maintenance: 2.0% isoflurane in air). Anesthetic depth and body temperature were monitored as above. The cameras were positioned to have a sharp image of the entire eye, with the mounting arms adjusted such that the cameras and mounting system caused minimal disruption to the mouse’s lateral and frontal field of view. Mounting arms were secured with cyanoacrylate adhesive glue (Histoacryl, B. Braun, Melsungen, Germany). The eye-camera system was then removed and the animal allowed to recover.

### Training procedure

Mice were acclimated to cricket capture in their home cage. Individual crickets were placed in the mouse’s home cage overnight, in addition to their standard *ad lib* mouse food. Mice were handled and habituated to the experimenter, the head cameras, and the head tracking mounts. The ability of each mouse to visually track the crickets was assessed using the protocol of Hoy et al.(Hoy, Yavorska et al. 2016). Briefly, the ability of the mice to track and capture crickets in a white walled, 480 x 375 x 210 cm arena was assessed in lit and dark conditions (Figure 7C). Mice were given 2 minutes in which to capture the crickets. Prior to the assessment mice were food deprived overnight before the trial.

### Placement of torsion tracking marks

Crenellations along the iridial-pupil border were less distinct in mice than those previous described in rats (Wallace, Greenberg et al. 2013). Ocular torsion changes were therefore measured by tracking the rotations of small spots of titanium dioxide (TiO_2_) paste dots (~ 300 μm) applied to ventral and/ or temporal locations on the cornea as described in (van Alphen, Winkelman et al. 2010). The TiO_2_ paste consisted of TiO_2_ powder (Kronos Titan GmBH, Leverkusen, Germany) mixed with a small quantity of sterile Ringer’s solution. Application of the TiO_2_ spots was performed with the animal anaesthetized with isoflurane (induction: 5% isoflurane, maintenance: 0.5-1.0% isoflurane in air, total time anesthetized 5-10mins). Anaesthetic depth and body temperature was monitored as above. Following application of TiO_2_ spots, mice were allowed to recover for >45 minutes prior to a cricket hunt. The presence of the TiO_2_ marks did not significantly change the animal’s cricket hunting performance as assessed by the average time taken to capture crickets (Figure 7C).

### Experiment procedure

Initially, mice were allowed to explore the experimental arena (1×1×0.26 m) without head camera mounts. During subsequent training sessions mice were acclimated to cricket hunting, with the head cameras on, in the experiment arena. Auditory white noise (60-65 dB, NCH-Tone generator v 3.26, NCH Software, Inc. Greenwood Village, USA) was provided through 4 speakers (Visaton, Germany), one on each wall of the arena. Olfactory noise was provided by ventilating the arena (TD-1000/200 Silent fan, S&P, Barcelona, Spain) through a perforated floor (5cm perforation spacing) with air blown through a cage containing live crickets (cricket cage dimensions 480×375×210cm). During experiments the arena was lit by a single lamp (4000 K, 9W, Osram, Munich, Germany) positioned ~1m above the arena. During each experiment the mouse was given 5 minutes to explore the arena without head cameras. After this period the mouse was removed from the arena and the head cameras were mounted. At the commencement of each hunt the cricket was released at a variable location into the central region of the arena.

### Eye camera and head position tracking system

Head and eye tracking was performed as described in (Wallace, Greenberg et al. 2013), with modifications as described below. The eye camera mount and implant were re-designed to be smaller, lighter and stronger (Figure 7A-B). The camera system body, camera holders and mounting arms were produced using a Formlabs Form2 SLA 3D printer (Formlabs Inc., USA), with Dental SG Resin (Formlabs Inc., USA) as the primary construction material. The cable used for data transfer and camera and position tracking LEDs power inputs was a flat flexible printed circuit (Axon Kabel GmbH, Leonberg, Germany). Eye movements were recorded at 60 Hz (camera resolution 752×480 pixels), with illumination provided by a ring of three IR-LEDs (λ=850 nm, OSRAM SFH4050 or SFH4053 @ 70mA, RS Components, Germany) surrounding the camera lens. The mouse’s head position and head rotations were tracked using seven IR-LEDs (λ = 950 nm, OSRAM SFH4043 @ 70mA, RS Components, Germany) mounted on three struts of carbon fiber that projected from the body of the camera system. The resultant total system weight was ~3g, including effective cable weight.

### Mouse head and cricket position tracking

The positions of the cricket within the arena were recorded using 4 cameras (488 x 648 px, recorded at 200 Hz, piA640-210gm, Basler cameras, Basler Ahrensburg, Germany) fitted with NIR-blocking filters (Calflex X, Qioptiq, Germany). Cameras were located ~1.5 m above the arena, and were positioned so that the arena was covered at all points by 2 or more cameras from differing vantage points. Mouse IR-head tracking LEDs were recorded at 200 Hz using 4 cameras (piA640-210gm, Basler cameras, Basler Ahrensburg, Germany). Image acquisition, synchronization and mouse head rotation calculations were performed as described previously (Wallace, Greenberg et al. 2013).

### Anatomical model

Head mount features and mouse anatomical features (medial canthi and nostril positions) were recorded at 50 Hz using four synchronized cameras (acA2040-90 um, Basler cameras, Basler Ahrensburg, Germany) fitted with 25 mm focal length objectives (CCTV lens, Kowa Optical Products Co. Ltd, Japan) calibrated as described for the overhead cameras in (Wallace, Greenberg et al. 2013). Cameras were positioned to provide images of the animal and headset from different angles to allow triangulation of the anatomical features (Figure 7D). During acquisition of the calibration images, the animal was illuminated with 12 IR-LED modules, (λ = 850, Oslon Black PowerStar IR-LED module, ILH-IO01-85ML-SC201_WIR200, i-led.co.uk, Berkshire, UK) run at 1A. Position tracking LED, medial canthi, nares, mirror corner and camera chip corner positions were marked in 2 or more camera views, in multiple synchronized frames. Based on the triangulated positions of anatomical features, head cameras and position tracking LEDs the mouse’s eye position could be placed a common coordinate system.

To establish the animal’s horizontal plane from the head tracking LEDs, a position for the animal’s nose was first defined by averaging to 3D positions of the marked nostrils. A pre-forward vector was calculated using the direction between mean of eyes and nose and a pre-up vector as vector orthogonal to the pre-forward and vector between the eyes. Next, the left vector was defined as orthogonal to pre-forward and pre-up. Finally, the system was rotated by 40° around the left vector such that forward vector was elevated. This established a head-fixed forward-left-up coordinate system that was based on the bregma-lambda sagittal plane by tilting the eyes-nose plane by an angle of 40°.

### Interpolation

Head tracking frame rates were 200Hz, while eye tracking cameras recorded at 60 Hz. Eye positions were consequently interpolated as follows: Let

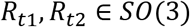

be two rotations that transform the vector (0,0, −1)^*t*^ into the gaze vectors *v*_*t*1_, *v*_*t*2_ in head fixed coordinates at times *t*1, *t*2. Then for a time *t*′ with

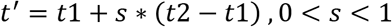

the corresponding rotation *R*_*t*′_ is interpolated such that *v*_*t*′_ is placed on the geodesic defined by *v*_*t*1_, *v*_*t*2_ with an angle of *s* * ∠ (*v*_*t*1_, *v*_*t*2_) to *v*_*t*1_, and the rotation of a vector perpendicular to (0,0, −1)^*t*^ is continuous and uniform between *t*1 and *t*2.

### Camera calibration

Overhead cameras for animal position and pose tracking, tracking of crickets and the cameras used for generation of the anatomical model were calibrated as described for the overhead cameras in (Wallace, Greenberg et al. 2013) and the eye camera calibration performed as described in (Wallace, Greenberg et al. 2013).

### Pupil position and pupil torsion tracking

Pupil boundary tracking, compensation for eye image displacement, and gaze vector calculation was performed as described previously in (Wallace, Greenberg et al. 2013). Where contrast between pupil and iris was insufficient to allow automated pupil position tracking, pupil positions were manually tracked.

The TiO_2_ spots for tracking ocular torsion were tracked manually in each image frame. Torsional rotations were determined based on the tracked TiO_2_ spot positions as follows. Total rotation of the eye was defined as previously described in (Wallace, Greenberg et al. 2013), as:

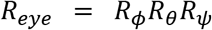

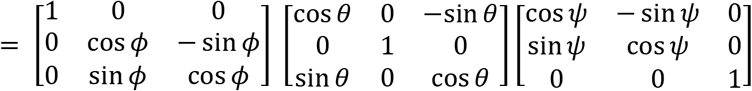

where *ϕ*=vertical, *θ*=horizontal and *ψ*=torsional rotations. The mouse’s gaze vector has the coordinates [0 0 −1]^*T*^ for the reference position of the eye, and in each frame:

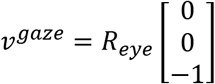

With the eye in its reference position, we assume that the marked TiO_2_ spot is located in the x-y plane of the eye camera (Wallace, Greenberg et al. 2013). The anatomical location of this marked spot can then be described by two unknown parameters *r* (where r>1 is the 3D distance of the eyeball surface to the eyeball center, and a distance of 1 describes the rotation radius of the pupil) and *α* is the fixed angle between the TiO_2_ mark and the gaze vector. After eye rotation the 3D location of the TiO_2_ is:

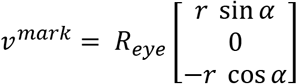

and the predicted pixel coordinates of the spot in the image are:

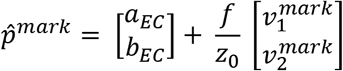

where *a_EC_* and *b_EC_* are the location in the image of the center of the eye ball and 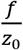 a scaling factor, both of which are determined in the calibration procedure for pupil boundary tracking, described in full in (Wallace, Greenberg et al. 2013).

When *r* and *α* are known the value *ψ* can be determined. Using the Matlab function **fminbnd** the squared 2D distance

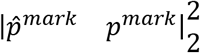

between the predicted and marked locations of the TiO_2_ mark is minimized.

This method is used to determine the ocular torsion based on the TiO_2_ spot location, both during and after calibration. Calibration was performed as follows:

For a given *r* and *α* choice, *ψ* can be calculated as above. The sum of square errors in pixel locations is then calculated over all frames. We optimized over *r* and *α* using the Matlab function **fminsearch**. To initialize *r*, we make use of the fact that the pupil model, *p^mark^* and *r* together determine the 3D location of the mark *v^mark^* in each image. For each frame we first calculated:

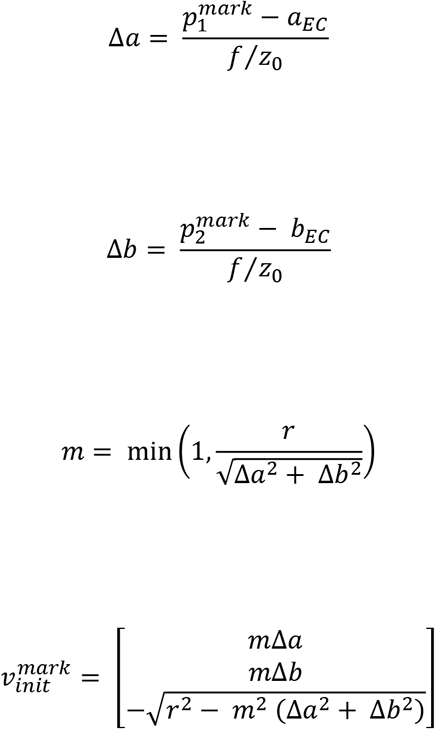

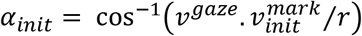

Using this method *α_init_* is estimated separately for each frame, and if the choice of *r* is correct then these values should agree. We can use **fminbind** to minimize the following with respect to *r*:

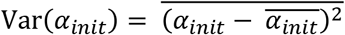

After *r* is initialized, *α_init_* is calculated, with *α* initialized using the mean over frames.

Torsional rotations were normalized by calculating a mean torsion value for the 0.01 % of frames that were closest to both median pitch and roll of the head. Torsional values in other tracked frames were then normalized to this mean torsion value.

### Cricket Position Tracking

Cricket body positions were automatically tracked using the method and algorithm described for tracking eye corners, as described in the section “*Compensation for lateral eyeball displacement – tracking of anatomical landmarks around the eye”* in (Wallace, Greenberg et al. 2013). To increase the contrast between the region around the cricket in the image and the cricket, ~100 background image frames (in which neither mouse nor cricket was present) were averaged and subtracted from frames in which the cricket was present. In frames where automated cricket position tracking was not possible, frames were tracked manually. As the cameras used for cricket tracking had been calibrated along with the animal position tracking cameras (see above), the 3-dimensional location of the cricket could be triangulated in a common coordinate system with the animal’s position.

### Classification of behavioral periods

To decrease the effects of tracking noise and rapid head rotations, mouse velocity, target bearing and inter-animal Euclidean distances were first filtered using a 50ms sliding window Gaussian filter.

The criteria used to classify the different hunt phases were based on those described in (Hoy, Yavorska et al. 2016). In an initial step, behavioral end points (**t**_end_) for capture periods were identified by manual inspection of the tracking movies. Further identification of the behavioral start points (**t**_start_) and **t**_end_ points for the different hunt sequence epochs were then identified as described below.

The **t**_end_ points were defined as:

A. The **t**_end_ point for a detect period was defined as the last frame before (1) Mouse head velocity in the direction of the cricket was >= 20 cm/s, (2) The mouse’s bearing towards the cricket was constantly below 90° and (3) the Euclidean distance between the mouse and cricket was continuously decreasing.
B. The **t**_end_ point for a tracking period was identified by locating local minima in the mouse-cricket Euclidean distance time plots, where local minima were defined as points at which the mouse came within a contact distance of 6 cm (measured from the tracked point on the mouse’s head, giving a > 3 cm separation between the mouse’s nose and the cricket). These were followed either by a capture period (see below) or were followed by a ⩾ 5cm increase in inter-animal Euclidean distance, which were defined as cricket escapes. In cases where the absolute value of the target bearing was > 90° before the mouse turned towards the prey, the start of the tracking period was taken as the first frame in which the bearing to the target was <90°. Only tracking periods, in which the initial Euclidean distance between the mouse and cricket was >20 cm were analyzed.
C. The **t**_end_ point for the capture period was taken to be the point 6 cm away from the cricket, following which a cricket captured and consumed.

The start points of the hunt epochs were defined as follows:

A. The **t**_start_ for the detect period was the frame 500 ms prior to the detect **t**_end_ point.
B. The **t**_start_ for the tracking period was the first frame after the **t**_end_ detect frame.
C. The **t**_start_ for the capture period was either; (1) the first period in which the mouse approached the cricket and directly caught it, or (2) the first frame in which the mouse approached the cricket and all subsequent cricket escapes (prior to the final cricket capture) were less than 5cm outside the contact zone (11 cm inter-animal Euclidean distance).

Cases in which the eye cameras were dislodged by the animal during the chase (n=4 hunt sequences) were included in the dataset up until the point where the cameras were dislodged.

### Target bearing

Target bearing was defined as the angle between the cricket position and the mouse’s forward head direction in the horizontal plane.

### Digital reconstruction of arena

For the digital reconstruction, the company 3dScanlab (Cologne, Germany) was engaged to create a complete scan, photo series and 3D mesh model of the arena and room, which they performed using an RTC 360 3D laser scanner (Leica, Germany). The 3D point cloud produced by the laser scanner was converted to a 3D mesh model, to which textures of the experiment arena obtained from photographs (Nikon D810, 36 mpx) were baked.

The camera tracking coordinate system, in which the mouse and cricket positions were tracked, and the scanned coordinate system of the 3D mesh model were aligned based on 16 fiducial points which could be clearly identified in both tracking camera images and the scan. Crickets were modelled as 2cm diameter, 1 cm thick disks centered on their tracked position with the disk’s axis oriented parallel to gravity.

### Generation of animal’s eye view

Each eye was modelled as a hemisphere with a 180° field of view whose equator was perpendicular to the animal’s gaze vector. For the projection of the environment onto the cornea, frame-wise animal’s eye views for both eyes were created with custom written software in C++ (g++ 7.5.0, QMake 3.1, Qt 5.9.5, libopenexr 2.2.0, libpng 1.6.34 and OpenGL-core-profile 4.6.0) on a GeForce RTX 2070 (NVidia driver 450.66), using first cube mapping followed by a transformation into a spherical coordinate system. To do this, individual frame-wise coordinate transformations were made using the eye locations and orientations determined as described above to transform the mesh model of the arena and cricket to a static eye coordinate system using custom written vertex shaders to perform the coordinate transformation and the fragment shaders to texture the mesh. A cube-map (1024 x 1024 pixels per face) was created by performing such coordinate transformations for a 90 degree view in the direction of the optical axis of the eye and four mutually orthogonal directions. Custom written code was then used to transform the cube-map into a spherical coordinate system, with a 180 degree opening angle, using vertex shaders, resulting in a 1024 x 1024 pixel frame exported as png and OpenEXR files. In addition to the color map, maps of depth (pixel-wise object intersection distance), object identification and 3D position of the object intersection point in the contralateral eye’s coordinate system were also generated.

### Prey image probability density maps

For generation of the prey image probability density maps, animal’s eye views were rendered that contained the cricket only (i.e. without inclusion of arena and room). Density maps from multiple detect-track sequences, and multiple animals, were made by averaging.

### Ocular Alignment

Ocular alignment was defined as the consistency of the projection of a given point in the eye view of one eye into the other in an infinitely distant environment. This is equivalent to a projection in an idealized finite-distant spherical environment while assuming a distance between the animal’s eyes of 0. For calculation, the radius of the sphere can then be set to 1 (without loss of generality). A point, located at the center of mass of the functional focus in each eye, was chosen from which to calculate the degree of inter-ocular alignment. This point was projected from one eye to the sphere surface and into the contralateral eye. The degree of alignment between the two eyes was calculated as follows:

Let

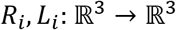

be the affine transformations for the left and right eye, and let

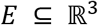

be the idealized environment. For a given direction *u* ∈ *S*^2^ we calculate the projection into the right eye *p_i_* ∈ ℝ^3^ by:

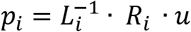

The average alignment is then calculated using the−formula:

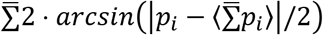

where 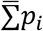 denotes mean and 〈 〉 denotes normalization.

### Visual field overlap

Visual field overlap was analyzed in the idealized finite-distant spherical environment described above for ocular alignment. Visual overlap was calculated from the frame-wise maps of 3D object intersection points in the contralateral eye (see above section “*Generation of animal’s eye view”*) generated for the ocular alignment analysis: pixels whose 3D object intersection points had an angle of less than 90° to the optical axis were considered part of the overlapping field of view. Probability maps of overlap were calculated by averaging.

For analyses of the effect of freezing eye movements, eye rotations (horizontal, vertical and torsional) were set to the mean rotation in one eye, and the effect quantified in the other eye view.

### Optic flow

To calculate the optic flow in a given pixel for a given eye, we consider the difference vector between the 3D positions in the static eye coordinate system of the object intersection point for this pixel one frame before and after the frame of interest, divided by 2 ∙ *dt* and mapped to unit distance by dividing by the distance between eye and interception point. This yields a 3D motion vector which is independent of influences of the frame rate. The spherical projection used in the rendering process described above is a non-conformal, locally non-isometric map, meaning that angles between lines and distances between points are not preserved. This makes it necessary to evaluate the flow in each point in a local, orthonormal 3D coordinate system defined by the direction vector between the eye position and the object intersection point and derivative vectors along the angular coordinates *vθ* and *vφ* at that point. Thus, we define the 2D flow at a given point as the orthogonal projection of the 3D flow vector onto the local plane spanned by *vθ* and *vφ*. In this study, we only use the first two components of the vector, while the third component contains the motion in radial direction to the eye.

In Figure 5C optic flow was calculated for the animal in the idealized spherical environment described above, meaning the animal’s head was equidistant to the surrounding at all points. This simplified scene was characterized as follows. Let

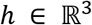

be the coordinate of the center of the mouse’s head, then the scene around it was defined as

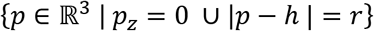

with r = 50 cm. For optic flow calculations the sphere is considered fixed in global coordinates, and the flow is evaluated at the point where the mouse is in the center of the sphere translating forward at a speed of 1 cm/s.

In Figure 5E optic flow was calculated with the animal in the digitally reconstructed environment (see above).

### Coloring of optic flow poles in mouse corneal views

The points in the scatter plot of optic flow poles in mouse corneal views were color-coded for the density of neighboring points using a two-dimensional Gaussian smoother with standard deviation

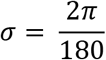

For a given point, the density was calculated as:

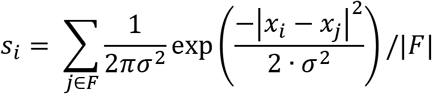

where *F* is the set of all considered frame indices, and

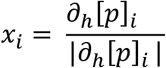

where *∂_h_*[*p*]_*i*_ is the discrete central difference quotient of the mouse’s eye trajectory *p* in frame *i,* in the coordinate system of the respective eye, evaluated over h=4 frames.

### Mouse Eye Model

When constructing the eye model, we took experimentally determined values from (Barathi, Boopathi et al. 2008) (see Table 1). While we recognize that this study employed a different strain of mice to the one used here, the methodology used provides estimates of physical and optical parameters measured under conditions closest to those relevant for the current study. Further, variation of these parameters was not found to change the model to an extent that would influence the conclusions drawn from analyses involving the eye model (see below). These values distinctly define the spatial shapes and positions of the refractive components of the model eye (Figure 3A), as well as refractive indices for all but the lens, *n_lens_*. We further assume a pupil radius of 594 μm, which is the mean of constricted and dilated mouse pupil sizes from (Pennesi, Lyubarsky et al. 1998). We define the focal point of a bundle of rays as the point with minimal least squares distance to the rays. To optimize the missing refractive index *n_lens_* ∶ Ω → *R*^+^ inside the lens body Ω ⊂ ℝ^3^, we first calculated two lens models and optimized them such that the focal point of 10000 rays emitted from an object at 10 cm distance on the optical axis lay on the retina. The first model, for optimization of the lens surface, was derived with optimal constant refractive index *n_c_* ∈ *R*^+^over the volume. The second model, for lens gradient optimization, was derived with a smooth transition of refractive index to the anterior and posterior lens boundary, ie. *n_b_* = 1.333 on *∂*Ω. We then used Poisson’s equation Δ *n_g_* = *c*, and optimized the strength of the gradient *c* ∈ *R*^+^. We assumed the final lens model as a linear combination of these two models:

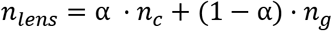

with *α* ∈ [0,1], where we optimized *α* as described for the above models, but from a point 10 cm away and 45° off optical axis. The derived refractive indices (Table 2) were within the range measured in (Cheng, Parreno et al. 2019).

To test the sensitivity of the model to changes in assumed physical parameters, we systematically changed the radius of curvatures listed in Table 1, and the thickness listed in Table 2 by 10, 50 and 100 μm (several different values were used, to check the linearity of the dependence). We calculated the propagation of uncertainty through the eye model by analyzing the variation of radial elevation on the retina of the 45 rays (above), taking the numerical differentiation of each input variable that was used in the model. Lens optimization was performed for each newly generated eye model (as described above). The maximum deviations were 0.4, 1.38, 2.76 degrees respectively for the 10, 50 and 100 μm changes (Figure 7E), and overall none of the observed effects on the model would influence the conclusions drawn from the analyses performed using the eye model.

### Projection from retina to cornea

The refractive elements in the rodent eye do not behave like ideally corrected optical elements, with the result that there is a distribution of incident rays with slightly varying angles of incidence on the cornea which converge on any given point on the retina. Projection from retina to cornea therefore requires an estimate of the distribution of outside world angles of incidence for any point of interest on the retina. To do this, we used a Monte-Carlo simulation to back-trace through the optics a set of randomly chosen rays emerging from the point of interest on the retina. Since the intensity of light on a surface with an incoming angle of θ is proportional to cos(*θ*), this function was also chosen for the probability density distribution of ray exit angles. The rays were then traced until they either hit any opaque surface, resulting in the affected ray being discarded, or passed through the anterior cornea, in which case the ray was accepted and its angle added to the distribution of passing exit angles for the respective point on the retina.

Refraction on boundary layers between different indices of refraction was performed analytically according to Snell’s law. In volumes with a continuous variable refractive index (i.e. gradient-index (GRIN) optics), we used a finite-elements model. We first discretized the lens as a 40×40×40 lattice of side length 2.4 mm. We then started from initial conditions where s(0) is the point of incidence and v(0) is the vector of incidence multiplied by the speed of light c. The subsequent discrete trajectory and direction of propagation is then calculated step-wise according to

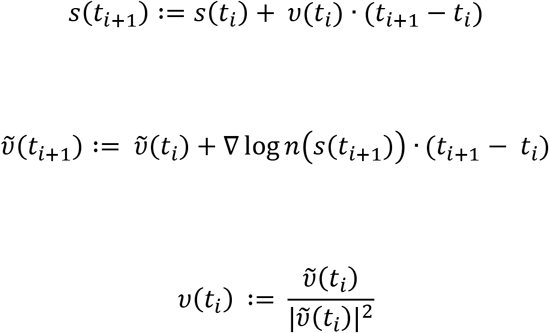

The gradient is calculated in the lens lattice as the three-dimensional difference quotient, and then linearly interpolated to the exact position *s*(*t_i_*) of the ray.

### Projection of retinal ganglion cell density contours onto the model eye cornea

To determine the corneal location corresponding to the histologically identified retinal specialization in the mouse, isodensity lines were redrawn from (Drager and Olsen 1980) in Illustrator and digitized using Matlab. Isodensity lines enclosing regions containing the highest and second highest density of retinal ganglion cells, as well as the optic disc and outline of the retinal whole mount, were redrawn directly from Figure 3A in (Drager and Olsen 1981), with horizontal being taken as horizontal (nasal-temporal) in the figure. The isodensity lines were scaled to match the eye diameter used for model eye, then placed into the model eye such that the center of mass of the optic disc reconstructed with the retinal ganglion cell contours was coincident with the intersection of the optic axis and retina in the eye model (Supplementary Figure 2A-C). As the eye model was rotationally symmetrical, no further alignment between the histology and eye model was necessary. The high retinal ganglion cell density regions were then back-projected from retina to cornea as described above (Supplementary Figure 2D-E).

### Eye in head coordinates

To quantify the effect of head rotations on VOR evoked eye movements in a common coordinate system, head rotations were normalized such that the average pitch and roll were 0. Axes were labeled X and Y respectively and eye rotations were represented using this horizon-aligned X-Y coordinate system. Positive head X values indicate head pitched up, while negative head X values indicate head pitched down. Negative head Y values indicate roll left, while positive Y values indicate roll right. Comparisons of the relationship between head and eye rotations were carried out using differential rotations between frame and average pose, defined in the following way: l′: L → G, r′: R → G, h: H → G are the affine transformations between Cartesian global coordinate system G, head-fixed coordinate system H and left/right-eye coordiante systems L/R.

The transformations from L/R respectively to H are:

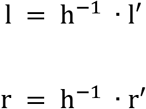

We calculate the left and right eye differential rotations as:

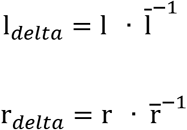

where 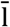 and 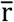 denote the average transformations over all frames (chordal L2 mean, implementation from SciPy 1.4.1).

### Statistical Analysis

Within one experimental trial, the experimentally measured variables of interest are highly correlated with each other. This fact prevents us from using standard statistical tests on the whole time-trace to establish if any difference we observed in the data across different experimental conditions are significant or not, as one requirement of these kind on tests is that the samples from the populations being compared are independent of each other. However, we realized that trial-to-trial variability is the dominant source of variability in the data, whereas within-trial variability explains a smaller fraction of the total variance observed (a more detailed report is found in Table 4). For this reason, we decided to represent each temporal trace by its median value. We used the median and not the mean, because the former is more resistant to the presence of outliers and it is better suited to represent the “average” value of a variable in this context. This operation reduced the size of the dataset to one data points per trial, which we can reasonably assume to be independent of each other.

---

**Movie 1.** Digitized and rendered view of the experiment arena and surrounding environment. Laser scanned and digitally reconstructed experiment environment, providing distance and positional information of objects within the mouse’s environment. When combined with the tracked 3D cricket positions and the tracked mouse head and eye positions and rotations this allowed the generation of a frame-by-frame mouse eye view of the prey and the surroundings.

**Movie 2.** Mouse eye views during cricket detection and tracking. Upper panels: Digitally rendered mouse left and right eye view’s of its prey (cricket - red) and the surrounding environment during prey detection, tracking and prey escape. Lower panel: recorded pursuit sequence. Green points indicate the tracked cricket body center. Note the transition from a peripheral monocular to a binocular lower nasal location within the visual fields. Note also the large overhead visual field.

## Author contributions

**1. Carl D. Holmgren**

Department of Behavior and Brain Organization, Research Center Caesar, Bonn, Germany

Contribution: Conceptualization; Methodology; Validation; Formal analysis; Investigation; Data curation; Writing – original draft preparation; Writing – review & editing; Visualization

Competing Interests: No competing interests declared

**2. Paul Stahr**

Department of Behavior and Brain Organization, Research Center Caesar, Bonn, Germany

Contribution: Methodology; Software; Validation; Formal analysis; Investigation; Data curation; Writing – review & editing; Visualization.

Competing Interests: No competing interests declared

**3. Damian J. Wallace**

Department of Behavior and Brain Organization, Research Center Caesar, Bonn, Germany

Contribution: Conceptualization; Methodology; Formal analysis; Data curation; Writing – original draft preparation; Writing – review & editing; Visualization; Supervision.

Competing Interests: No competing interests declared

**4. Kay-Michael Voit**

Department of Behavior and Brain Organization, Research Center Caesar, Bonn, Germany

Contribution: Methodology; Software; Validation; Formal analysis; Resources; Data curation; Writing – review & editing; Supervision.

Competing Interests: No competing interests declared

**5. Emily J. Matheson**

Department of Behavior and Brain Organization, Research Center Caesar, Bonn, Germany

Contribution: Methodology; Resources

Competing Interests: No competing interests declared

**6. Juergen Sawinski**

Department of Behavior and Brain Organization, Research Center Caesar, Bonn, Germany

Contribution: Methodology; Resources.

Competing Interests: No competing interests declared

**7. Giacomo Bassetto**

Department of Behavior and Brain Organization, Research Center Caesar, Bonn, Germany

Contribution: Formal analysis.

Competing Interests: No competing interests declared

**8. Jason N. D. Kerr**

Department of Behavior and Brain Organization, Research Center Caesar, Bonn, Germany

Contribution: Conceptualization; Methodology; Formal analysis; Data curation; Writing – original draft preparation; Writing – review & editing; Visualization; Supervision; Funding acquisition.

Competing Interests: No competing interests declared

## Funding

### Stiftung caesar and Max Planck Society

Carl D. Holmgren

Paul Stahr

Damian J. Wallace

Kay-Michael Voit

Emily J. Matheson

Juergen Sawinski

Giacomo Bassetto

Jason N. D. Kerr

The funders had no role in study design, data collection and interpretation, or the decision to submit the work for publication.

---

**Figure 1 – figure supplement 1.**
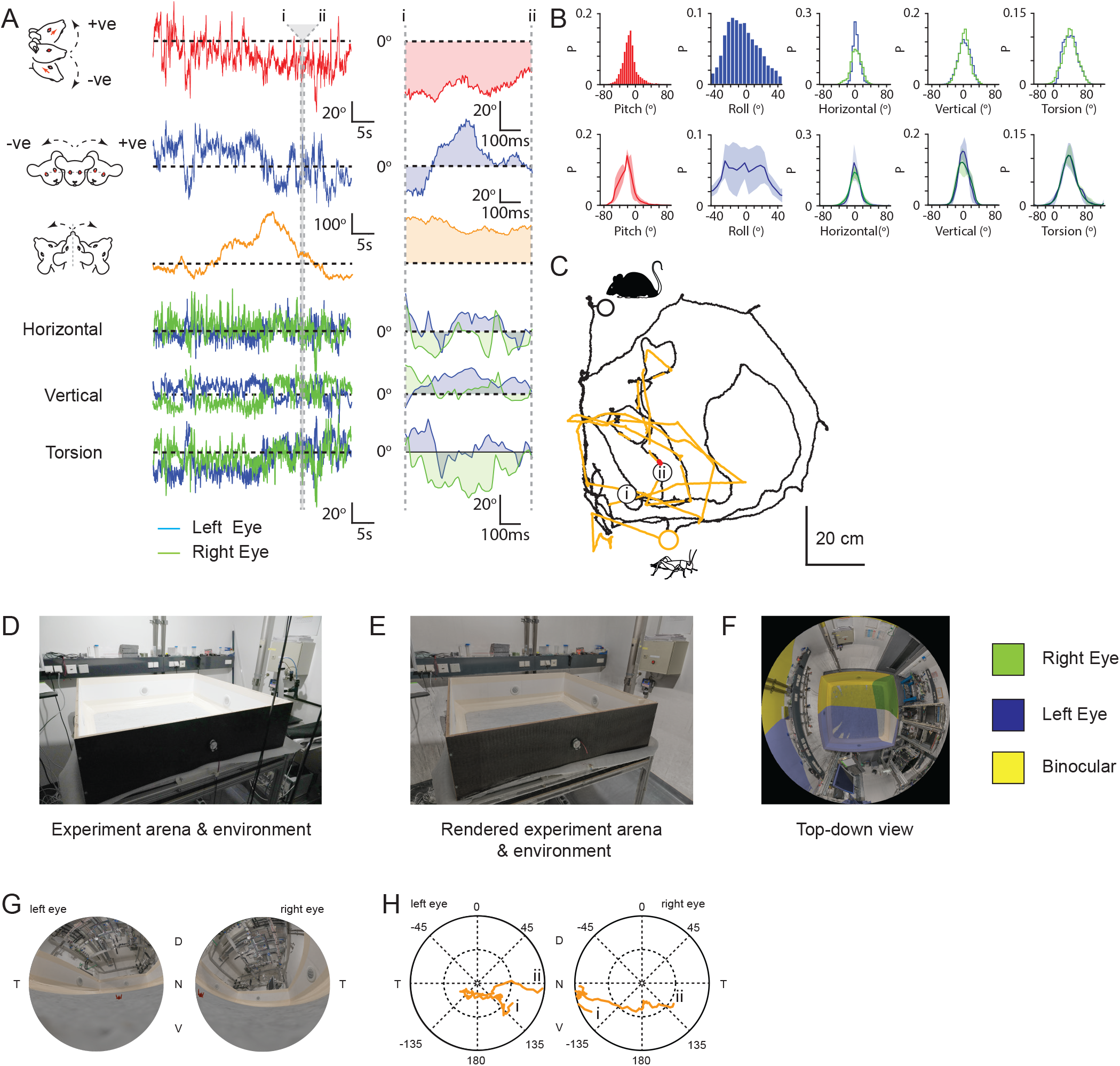
Generation of mouse eye views during cricket pursuit. **(A)** Head pitch (red), roll (blue) and yaw (orange) and associated left (blue) and right (green) horizontal, vertical and torsional eye movements during the 46.2s, example cricket pursuit sequence shown in C. (Right) Head and eye rotations during the 0.65s region between i and ii in the cricket pursuit sequence in C. **(B)** Example (upper rows) head pitch (547118 frames), roll (547118 frames), and horizontal (612161 frames), vertical (547118 frames) and torsional (612161 frames), eye rotations (n=1 animal). Lower rows: head and eye rotations from 3 mice. Data for B (lower), from 1436204 frames, from 3 animals. **(C)** Mouse (black) and cricket (orange) paths during a 46.2s segment of a single pursuit sequence for one animal. **(D)** Photograph of experiment arena and surrounding environment. **(E)** Digital rendering of the same experiment arena and surrounding environment. **(F)** Top-down view of the mouse’s left and right monocular and binocular fields of view (mouse’s head would be centered at the intersection point of monocular and binocular fields of view). **(G)** Cricket (red) position in the rendered left and right eye corneal fields of view of the experiment arena and surrounding environment during the pursuit sequence in C. **(H)** Trajectory of the projected cricket position in the left and right corneal views, during the pursuit sequence in C.

**Figure 2 – figure supplement 1.**
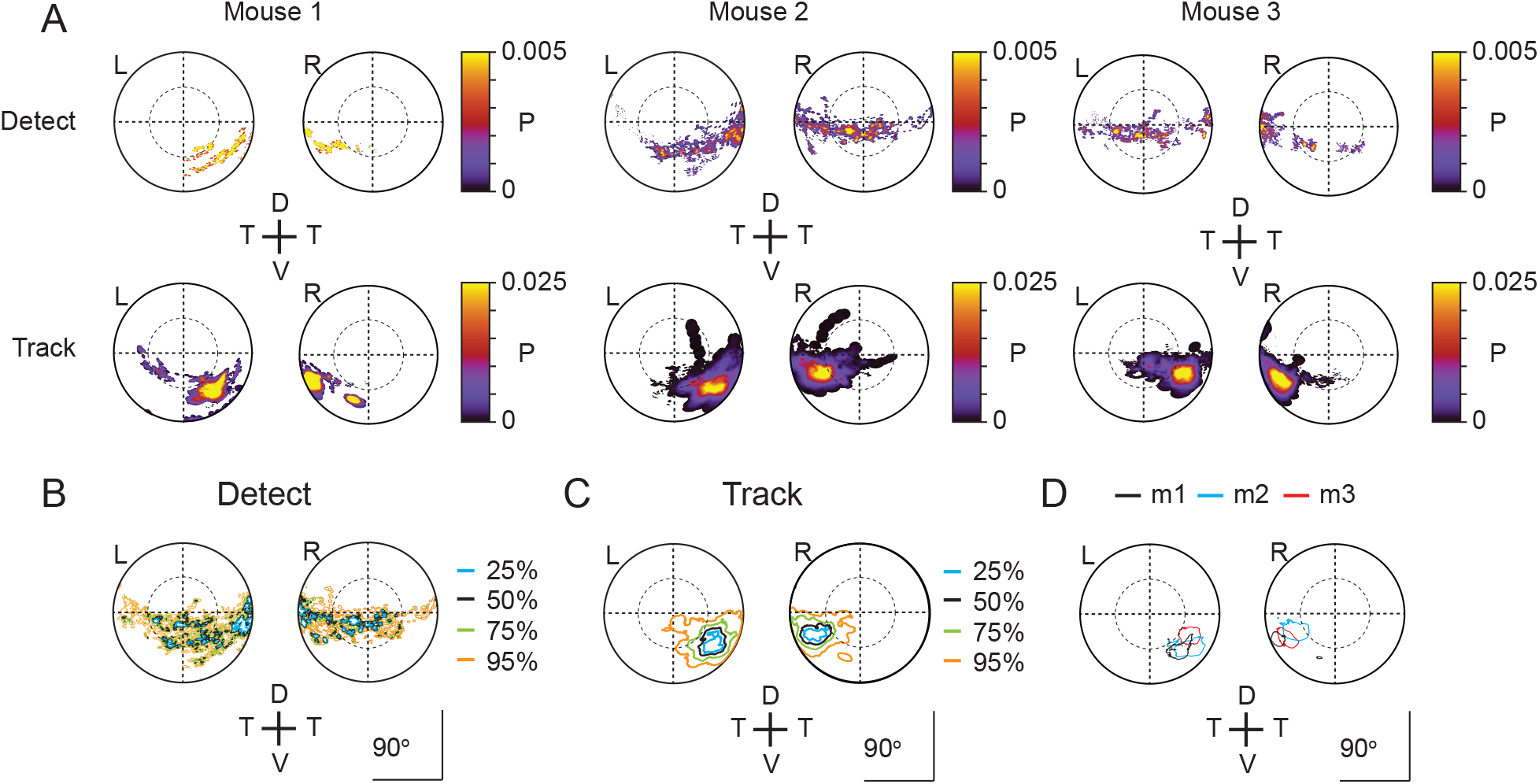
Individual corneal prey image heatmaps. (A) Probability density maps for detect (upper row) and track (lower row) epochs for each of the three animals individually. Data from 4 detect and 5 track sequences, 27 detect and 28 track sequences and 17 detect and 19 track sequences for mouse 1, 2 and 3 respectively. (B) Isodensity contours calculated from the average probability density maps for all detect epochs from all 3 animals. (C) Isodensity contours for all track epochs from all 3 animals. (D) 50% isodensity contour (defined as in Figure 2H) during track epochs for each of the three mice (m1-m3) individually.

**Figure 3 – figure supplement 1.**
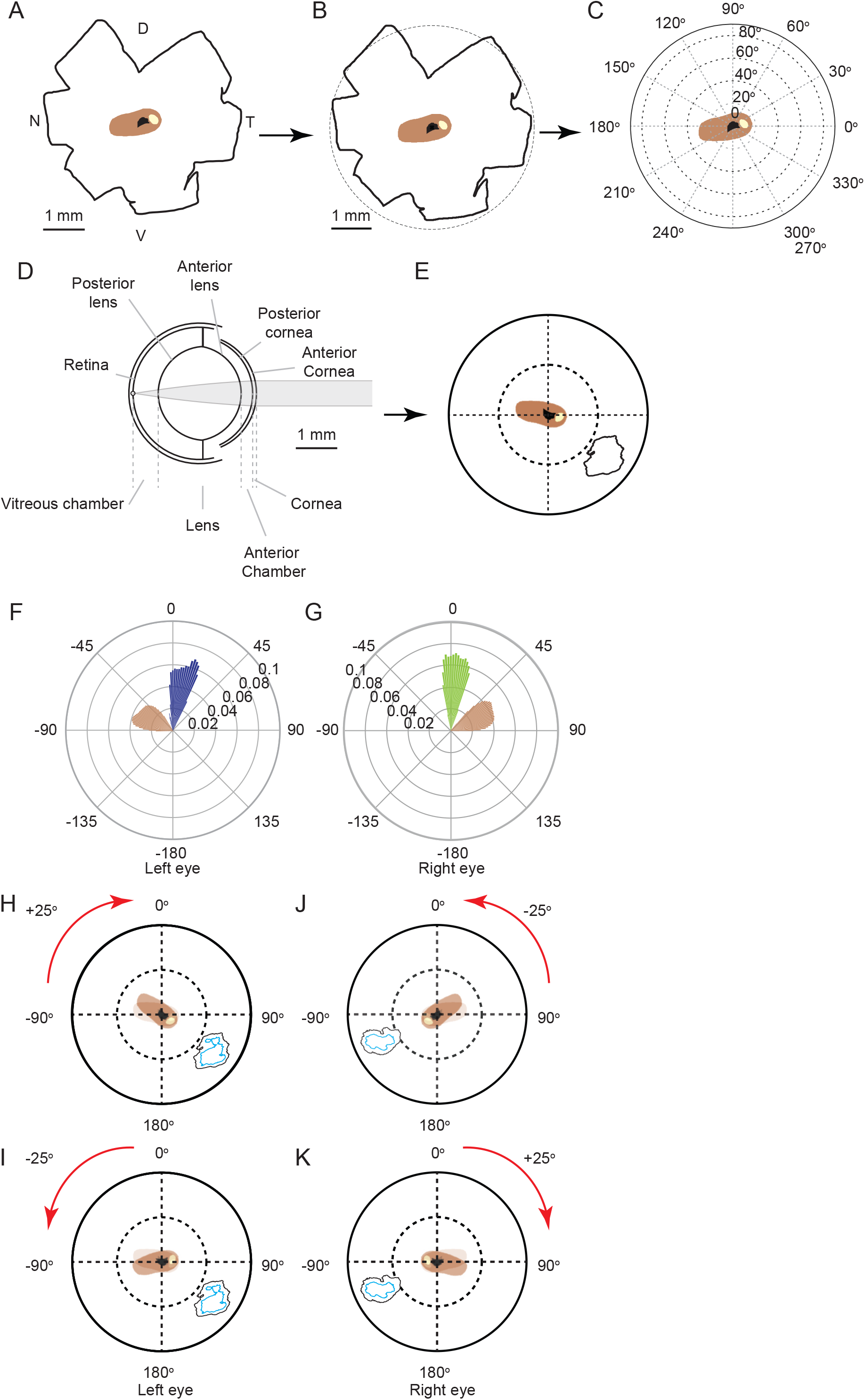
Projecting high retinal ganglion cell density region from retina to cornea. **(A)** Retinal whole mount redrawn from (Drager and Olsen 1981) including whole mount outline (black), and outlines of the optic disc (black) and highest (>8000 cells/mm2,beige) and second highest (>7000 cells/mm2brown) retinal ganglion cell density isodensity lines. **(B)** Overlay of the redrawn retinal whole mount from A and a representation of the mouse eye equatorial diameter (dashed) from (Tkachenko 2010). The center of the equatorial diameter was overlaid with the center of mass of the outline of the optic disc of the redrawn whole mount (black cross). Color coding as in A. **(C)** Retinal isodensity lines represented in spherical coordinates. Color coding as in A. **(D)** Schematic of mouse eye model (from Figure 3A). **(E)** Regions within the isodensity contours from A and the 50% isodensity contour from the track epochs from Figure 2H projected through the mouse eye model into the corneal view from the left eye (from Figure 3B). **(F)** Top-down view of the coverage region for the left eye of the 50% isodensity contour (blue) and second highest RGC region (brown). Bars represent the probability density function for the respective regions at that azimuth angle. Mouse’s forward direction directed to 0o, and mouse’s right directed to 90 o. **(G)** Top-down view of the coverage region for the right eye of the 50% isodensity contour (green) and second highest RGC region (brown). Conventions as in F. **(H & I**) left and **(J & K)** right eye corneal views, showing the effect on the orientation and location of RGC regions and isodensity contours of ± 25o torsional offsets. Original position of RGC region, beige, position after offset brown; color-coding of isodensity contours as in Figure 2H.

**Figure 4 – figure supplement 1.**
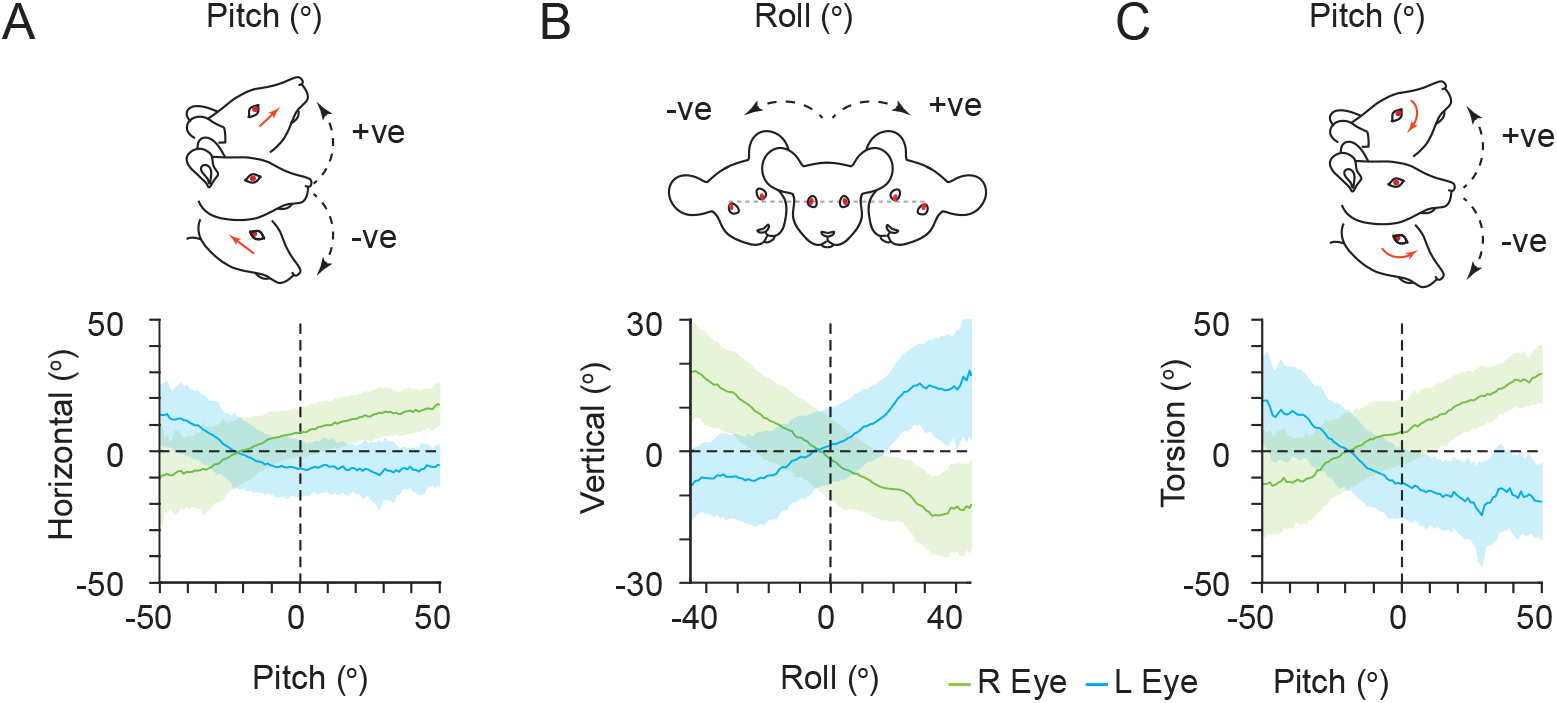
VOR relationships between head and eye rotations and alignment of left and right eyes. **(A)** Relationship between mouse head pitch and horizontal eye rotations (left eye, blue; right eye, green; mean±SD). **(B)** Relationship between head roll and vertical eye rotations. Plot conventions as in A. **(C)** Relationship between head pitch and torsional eye rotations. Plot conventions as in A. Data for A-C, from 1436204 frames, from 3 animals.

**Figure 4 – figure supplement 2.**
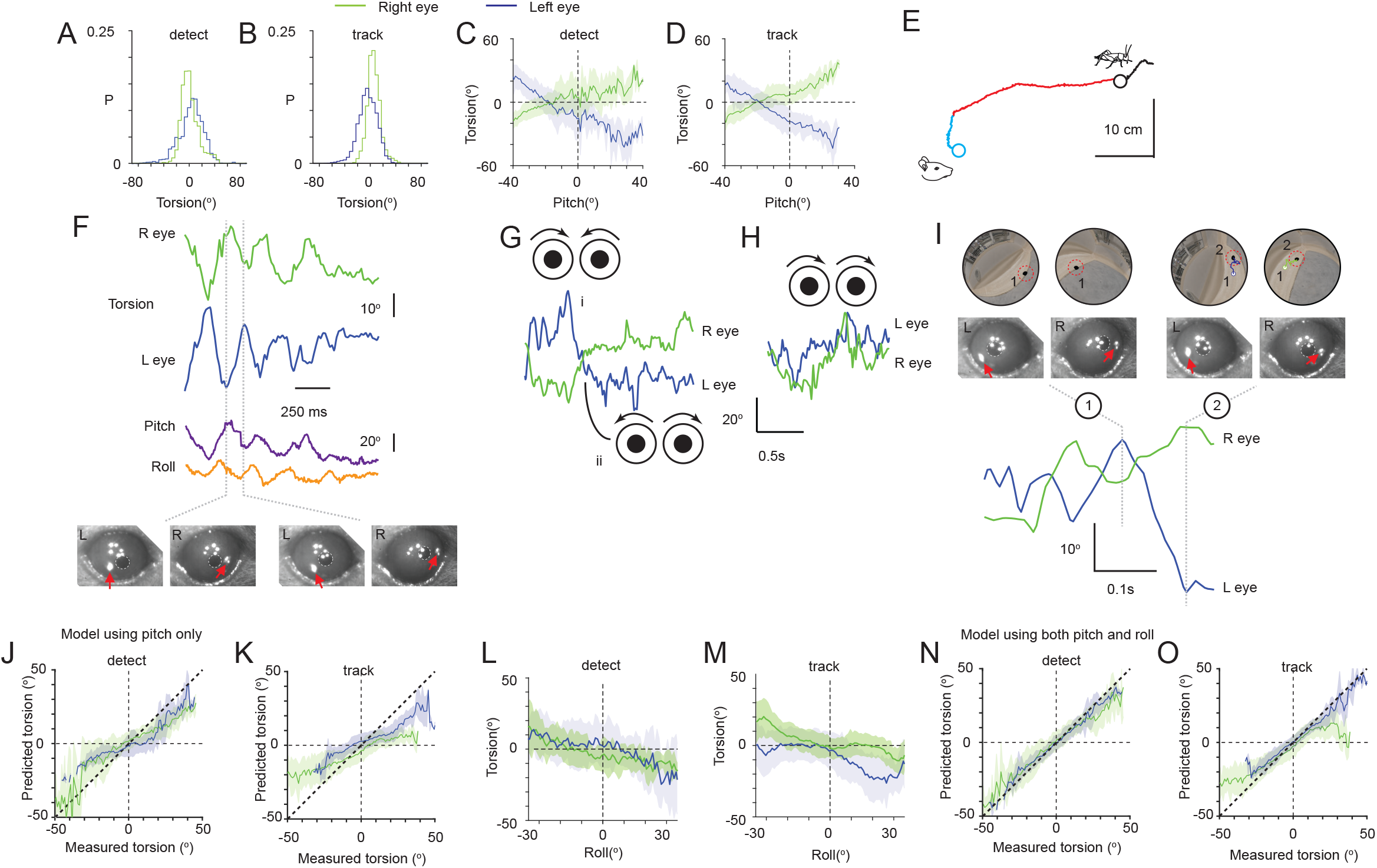
Ocular torsion during cricket pursuit. **(A)** Distribution of left (blue) and right (green) eye torsional rotations during detect epochs. Data from 57 epochs (4406 frames) from 3 animals. **(B)** Distribution of ocular torsion during track epochs. Conventions as in A. Data from 65 epochs (13624 frames) from 3 animals. **(C)** Average relationship (mean±SD) between head pitch and torsional eye rotations during detect epochs for left (blue and right (green) eyes. Data from 57 epochs (4406 frames) from 3 animals. **(D)** Average head pitch and torsional eye rotations relationships during track epochs. Conventions as in C. Data from 65 epochs (13624 frames) from 3 animals. **(E)** Mouse (detect epoch, blue; track epoch, red) and cricket (black) trajectories during one example pursuit sequence. **(F)** Torsional rotations of the left (blue) and right (green) eyes, and head pitch (purple) and roll (orange), during the pursuit sequence in E. Lower panels show example eye images from the indicated time points in the kinetic traces. Red arrows indicate tracked TiO_2_ spots. **(G)** Example sequences showing torsional rotation kinetic traces for left (blue) and right (green) eyes during in- (i) and excyclovergence (ii) from one pursuit sequence. Schematics show the ocular rotations in the left and right eyes. **(H)** Example sequence showing dextrocycloversion in one pursuit sequence. Conventions as in G. **(I)** Example of the effect of torsional rotations on prey image location. Corneal eye views of the cricket (black ellipse in red dashed circle) and arena (upper) and associated eye images (middle) at the time points indicated in the torsion kinetic traces (lower) for the left (blue) and right (green) eyes. Note cricket trajectories in left and right corneal eye views, which show the trajectory of the cricket in the corneal views between time points 1 and 2. Red arrows in eye images show TiO2 torsion tracking spots. **(J)** Performance of a model predicting torsion based on head pitch alone for left (blue) and right (green) eyes during detect and **(K)** track epochs. **(L)** Average (mean±SD) relationship between head roll and torsional eye rotations during detect epochs for left (blue) and right (green) eyes. Data from 57 epochs (4406 frames) from 3 animals. **(M)** Average head roll and torsional eye rotation relationship during track epochs. Conventions as in L. Data from 65 epochs (13624 frames) from 3 animals. **(N)** Performance of a model predicting torsion based on both head pitch and roll. Conventions as in J. For both J and N, data taken from 57 detect epochs (4406 frames), from 3 animals. **(O)** Performance of a model predicting torsion based on both head pitch and roll during tracking phases. For both K and O, data taken from 65 prey tracking epochs (13624 frames), from 3 animals.

**Figure 4 – figure supplement 3.**
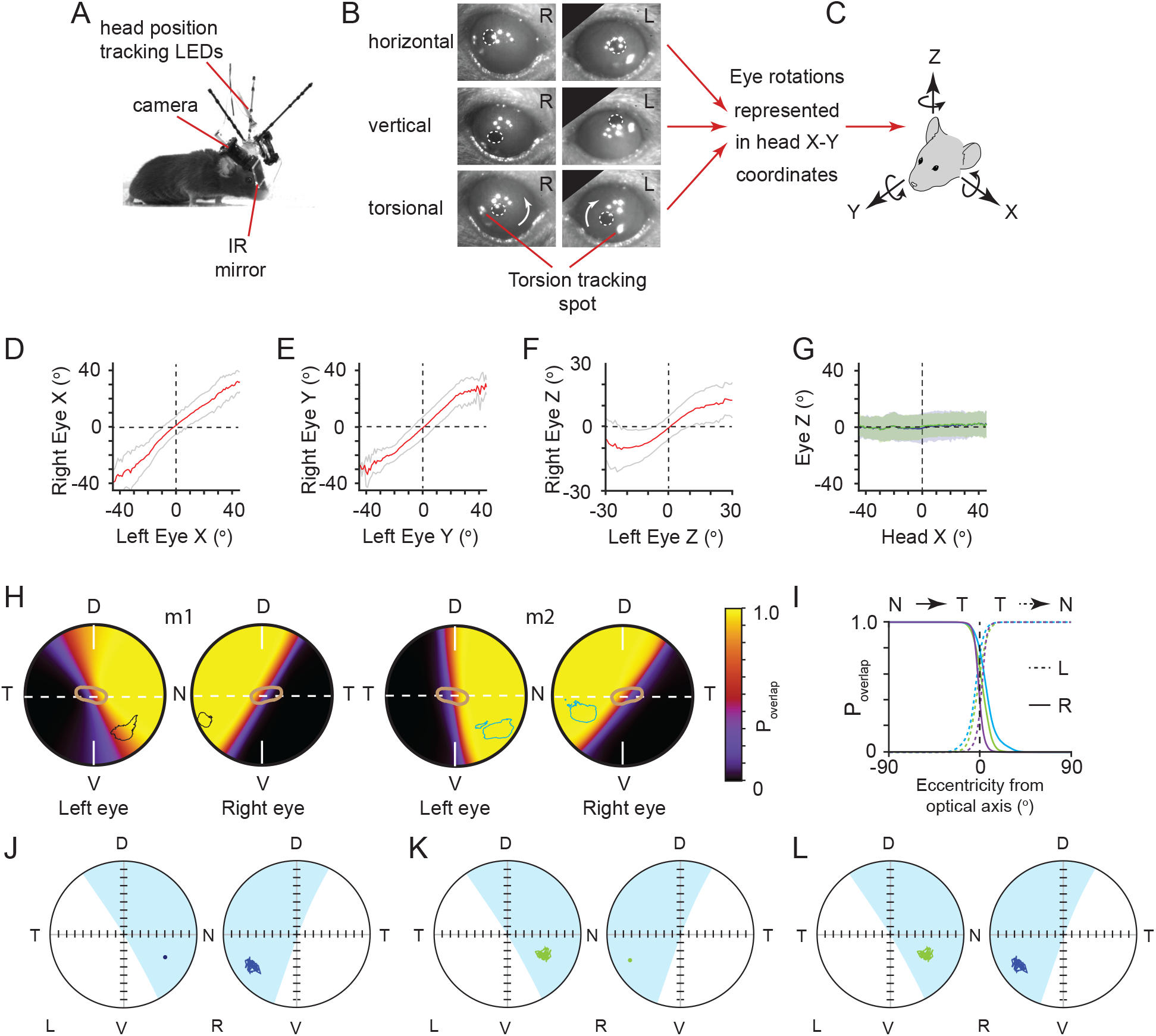
VOR relationships between head and eye rotations and alignment of left and right eyes. **(A)** Image of mouse with detachable miniaturized eye cameras and head position tracking system. **(B)** Example eye images showing horizontal, vertical and torsional eye rotations. Note TiO_2_ spots on the corneal surface for tracking torsion highlighted in lower panels. **(C)** Schematic of the common head and eye rotational axes. Relationship between **(D)** left and right eye X-rotations, **(E)** Y-rotations and **(F)** Z-rotations in common rotational axes. **(G)** Relationship between head X rotations and eye Z rotations for left eye (blue) and right eye (green). Data for D-G are represented as mean±SD, and are from 154500 frames from 3 animals. **(H)** Corneal view showing probability of overlap of left and right visual fields for two example animals m1 (left, 36449 frames) and m2 (right, 50874 frames), with overlay of isodensity contours (m1 -black, m2 - blue) from functional foci (see Figure 2 – figure supplement 1D) and contour of second highest RGC region (brown) from Figure 3B. **(I)** Profile of probability of overlap for left (dotted) and right (solid) eyes as a function of angular distance from optical axis for all three animals. Profile taken from horizontal axis through optical axis as shown in Figure 4D (dotted line in 4D, N = 3 animals, green = 36449 frames, blue = 50874 frames, purple = 71995 frames, respectively). **(J)** example of ocular alignment for the reference spot in the left eye projected into the right eye. **(K)** reference spot in the right eye projected into the left eye. **(L)** alignment over time for both reference spots.

**Figure 5 – figure supplement 1.**
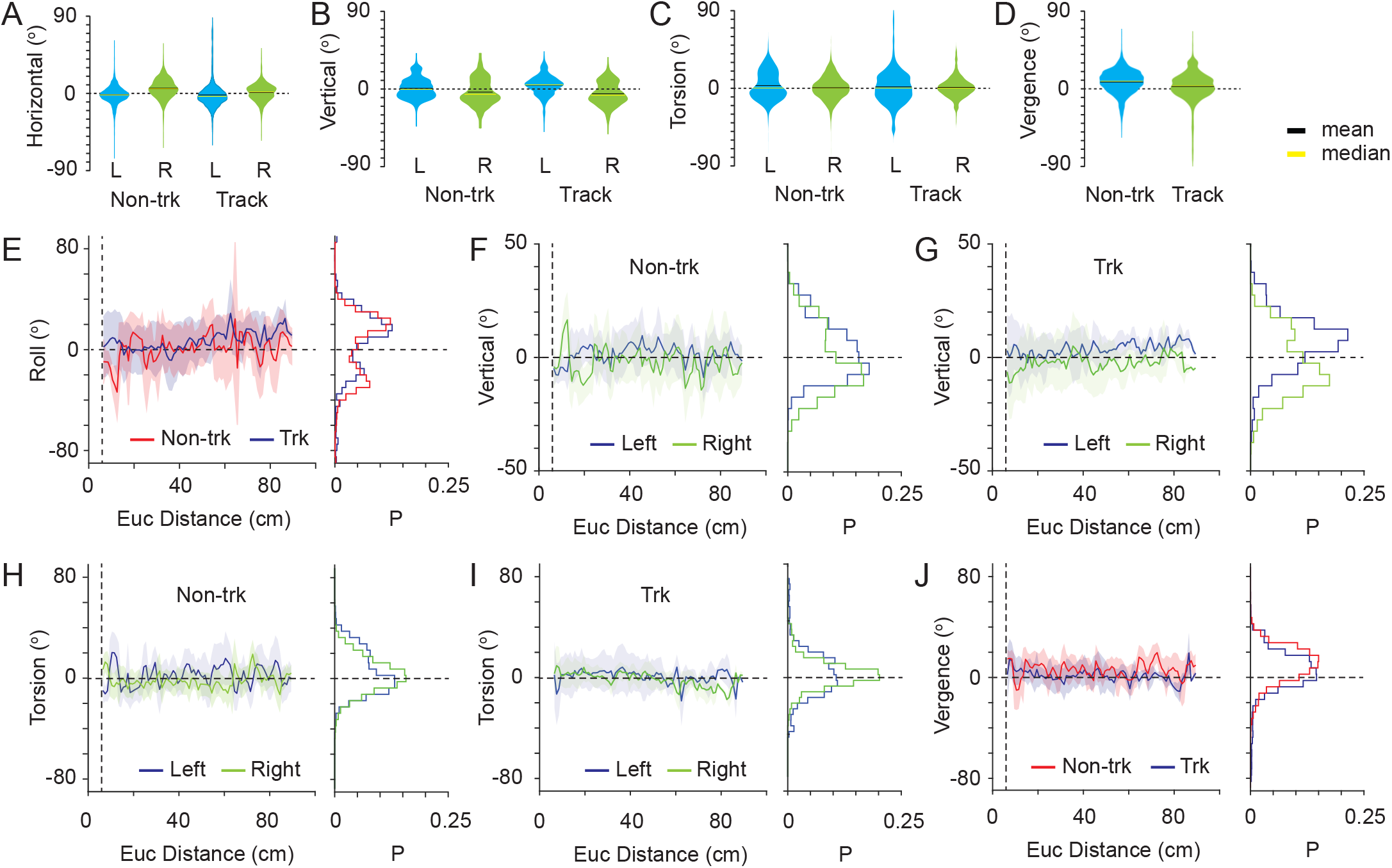
Eye movements during non-tracking and tracking periods. **(A)** Violin plots showing the variability in horizontal eye rotations for left (blue) and right (green) eyes during non-tracking (Non-trk) and track (Track) epochs. **(B)** Variability in vertical eye rotations during non-tracking and track epochs. Conventions as in A. **(C)** Variability in torsional eye rotations during non-tracking and track epochs. Conventions as in A. **(D)** Variability in ocular vergence during non-tracking and track epochs. Conventions as in A. **(E)** Average relationship (mean±SD) between head roll and Euclidean distance from mouse to cricket during non-track (red) and track (blue) epochs. Data histogram shown at right. **(F)** Average relationship (mean±SD) between vertical eye rotations of left (blue) and right (green) eyes and Euclidean distance between mouse and cricket during non-track epochs. Data histogram shown at right. **(G)** Average relationship between vertical eye rotations and mouse-cricket Euclidean distance during track epochs. Conventions as in F. **(H)** Average relationship between torsional eye rotations and mouse-cricket Euclidean distance during non-track epochs. Conventions as in F. **(I)** Average relationship between torsional eye rotations and mouse-cricket Euclidean distance during non-track epochs. Conventions as in F. **(J)** Average relationship between ocular vergence and mouse-cricket Euclidean distance during non-tracking and tracking epochs. Conventions as in E. For all panels, data taken from 18 non-track epochs (15649 frames) and 18 track epochs (8510 frames), from 3 animals.

